# Genomic characterization of world’s longest selection experiment in mouse reveals the complexity of polygenic traits

**DOI:** 10.1101/2021.05.28.446207

**Authors:** Sergio E. Palma-Vera, Henry Reyer, Martina Langhammer, Norbert Reinsch, Lorena Derežanin, Jörns Fickel, Saber Qanbari, Joachim Weitzel, Sören Franzenburg, Georg Hemmrich-Stanisak, Jennifer Schön

## Abstract

A unique set of mouse outbred lines has been generated through selective breeding in the longest selection experiment ever conducted on mice. Over the course of >140 generations, selection on the control line has given rise to two extremely fertile lines (>20 pups per litter each), two giant growth lines (one lean, one obese) and one long-distance running line. Genomic analysis revealed line-specific patterns of genetic variation among lines and high levels of homozygosity within lines as a result of long-term intensive selection, genetic drift and isolation. Detection of line-specific patterns of genetic differentiation and structural variation revealed multiple candidate genes behind the improvement of the selected traits. We conclude that the genomes of these lines are rich in beneficial alleles for the respective selected traits and represent an invaluable resource for unraveling the polygenic basis of fertility, obesity, muscle growth and endurance fitness.

## Introduction

Artificial selection is the selective breeding of organisms by which desired phenotypic traits evolve in a population^1^. Farm animals are the result of this selective breeding process to achieve efficient food production. However, artificial selection can also be applied experimentally in other species in order to connect genes and other genomic elements to selection response for complex traits such as behavior^2^ and limb elongation^3^.

The worldwide longest selection experiment on mice began in the early 1970’s at the former Forschungszentrum für Tierproduktion (FZT), nowadays called Leibniz Institute for Farm Animal Biology (FBN) located in Dummerstorf, Germany^4,5^. Starting from a single founder line, selection lines for different complex traits were bred with population sizes of 60-100 breeding pairs per line. An unselected control line from the same founder line was maintained over the entire selection period with a larger population size (125-200 breeding pairs)^4,5^. Over the course of >140 generations, selection has shaped the genomes of the Dummerstorf trait-selected mouse lines, leading to extreme phenotypes that include increased litter size (more than double the litter size of the unselected mouse line)^6^, body mass (approx. 90g body weight at 6 weeks of age)^7^ and endurance (up to 3× higher untrained running capacity)^8^.

In contrast to most murine models that mainly rely on targeted genetic modifications^9,10^, the Dummerstorf trait-selected mouse lines offer the possibility to elucidate the unpredictable genetic background of different complex traits, where multiple genes, regulatory elements and pathways act in conjunction to shape the phenotype the lines were selected for.

Here we describe the selection history of this unique selection experiment, characterize specific patterns of genetic variation and identify genes that are likely associated to each selected trait.

## Materials and Methods

### Selection history of the Dummerstorf trait-selected mouse lines

The selection experiment started in 1969 (Table 1 and 2, for more detail see Supplementary Material) with the establishment of a founder line FZTDU (<u>F</u>orschungs<u>z</u>entrum für <u>T</u>ierproduktion <u>Du</u>mmerstorf)^4,5^ by systematic crossing of four outbred strains (NMRI orig., Han:NMRI, CFW, CF1) and four inbred strains (CBA/Bln, AB/Bln, C57BL/Bln, XVII/Bln). From FZTDU five lines were established through selective breeding: two lines were selected for increased litter size (DUK and DUC), one for increased body mass (DU6), and one each for protein mass (DU6P) and treadmill running endurance (DUhLB) (Table 2, Fig. 1).

**Table 1.** Summary selection history of the Dummerstorf mouse lines.

|  | FZTDU | DUK | DUC | DU6 | DU6P | DUhLB |
| --- | --- | --- | --- | --- | --- | --- |
| Established (year) | 1969 | 1971 | 1971 | 1975 | 1975 | 1982 |
| No. Founders (BPs) | NR | 60 | 60 | 80 | 80 | 100 |
| Trait increment | -- | 2× | 2× | 3× | 2× | 3× |
| Percentage selected<sup>1</sup> | -- | 25-80 | 25-80 | 45-90 | 45-70 | 40-100 |
| Relocation<sup>2</sup> at generation | 160-164 | 165 | 163-164 | 154-155 | 154-155 | 120-121 |
| No. BPs per generation before relocation | 200 | 60-100 | 60-100 | 60-80 | 60-80 | 60-100 |
| No. BPs after relocation (founders) | 55 | 19 | 24 | 7 | 19 | 22 |
| No. BPs per generation (current) | 125 | 60 | 60 | 60-120 | 60 | 60 |
| End of selection (at generation) | -- | Ongoing | Ongoing | Ongoing | 152 | 141 |
| No. generations under selection<sup>3</sup> | -- | 182,189 | 180,187 | 169,177 | 152,152 | 117,117 |
| WGS at generation(s) | 188,195 | 188,195 | 186,193 | 177,185 | 177,184 | 143,150 |

BPs: breeding pairs. WGS: whole genome sequencing; NR: no records.
Trait increment: Mean trait expression in the sampled generation compared with trait expression in starting generation.
<sup>1</sup>Percentage selected = (number of selected/total number of animals per generation)×100
<sup>2</sup>Transfer of animals to a new housing building in 2011.
<sup>3</sup>Total generations under selection until first and second sampling

**Table 2.** Selection criteria for Dummerstorf trait-selected mouse lines.

| Line-ID | Selected Sex | Trait |
| --- | --- | --- |
| FZTDU | -- | Unselected |
| DUK | Females | Number of offspring in first litter and litter weight at birth |
| DUC | Females | Number of offspring in first litter and litter weight at birth |
| DU6 | Males | Body mass at day 42 of age |
| DU6P | Males | Protein amount in carcass at day 42 of age |
| DUhLB | Males | Submaximal untrained running distance in treadmill |

**Fig. 1.**
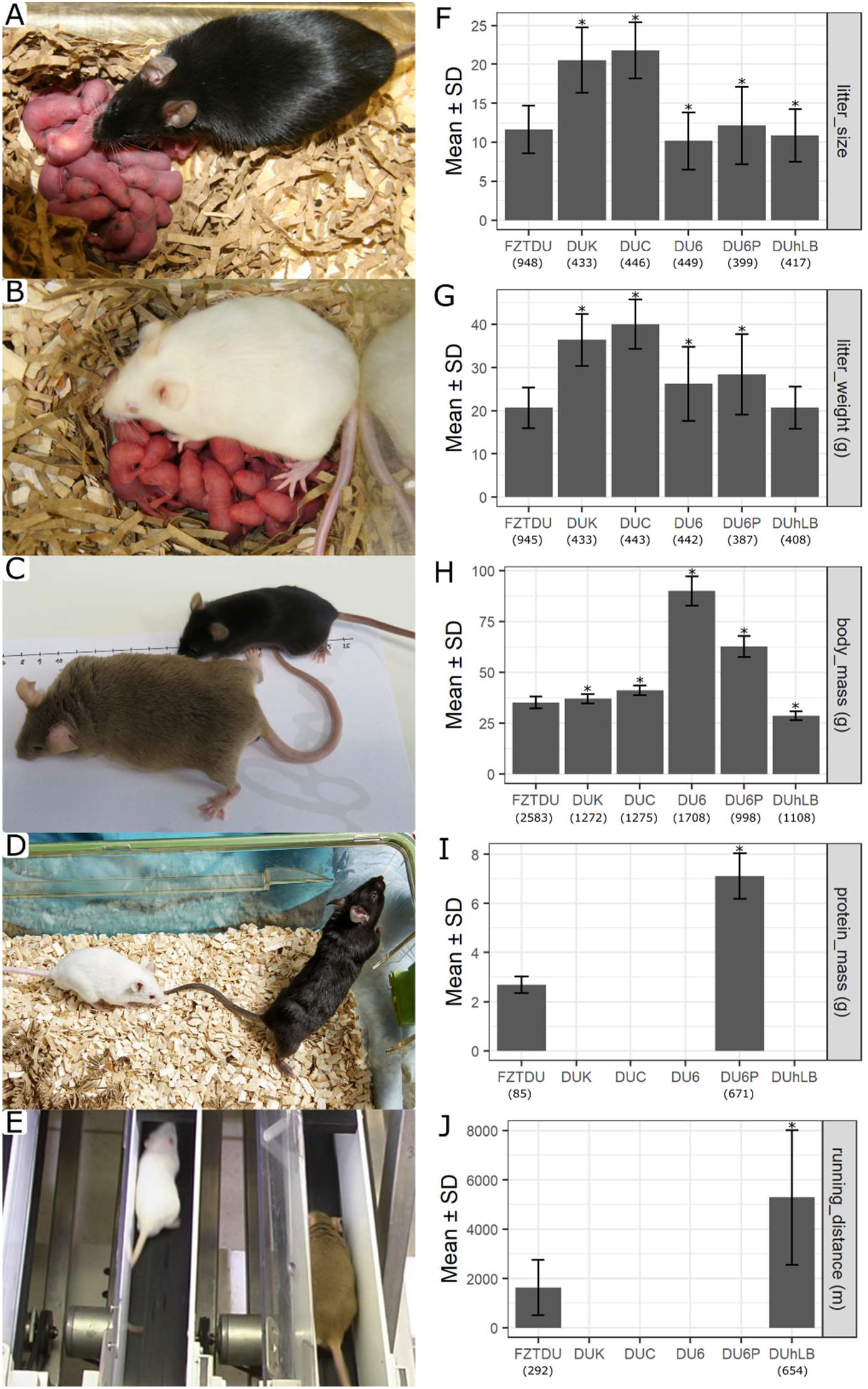
Phenotypic characteristics of the five trait-selected Dummerstorf mouse lines and the unselected control line FZTDU. Representative subjects showing the impressive litter size of DUK and DUC (A,B,F,G) and the considerable body size difference at six weeks of age between DU6 (C,H) or DU6P (D,H,I) relative to FZTDU. (E) Untrained mice undergoing a treadmill running endurance trial and the increased running performance of DUhLB due to selection (J). Stars signify differences (p < 0.05) after conducting a t-test between trait-selected lines and FZTDU. Sample sizes are indicated below tick labels (x-axis).

In general, all lines were developed through among-family selection^11^, whereby litters were ranked according to each trait of interest (Table 2) and then parents were chosen at random from the highest ranked litters. DUK and DUC were ranked by the total number of pups and by the total litter weight. However, for an interim period of 10 generations families were ranked only by litter weight. DU6 was selected for growth by ranking litters by the total weight of two randomly samples males from each litter at 42 days of age. Whenever possible these males were not chosen as sires. Similarly, the protein mass line DU6P was developed according to protein mass by combining weight and protein content of the carcass of a single male from each litter at 42 days of age. Occasionally protein mass could not be determined (e.g. because of technical issues or limited lab capacities), in which case litters were ranked by the combined weight of two males, as described for line DU6. Finally, the sire of each litter in line DUhTP was submitted to a treadmill test after the mating period and litters got their rank according to performance of their sires.

FZTDU was kept as an unselected control line with 200 breeding pairs, whereas the trait-selected lines were maintained with 60-100 breeding pairs. However, when animals had to be relocated to a new animal housing building in 2011 after 120-165 generations, the number of breeding pairs drastically dropped to 55 for FZTDU, ~20 for DUK, DUC, DU6P and DUhLB, and to as low as 7 for DU6 (Table 1). This transition could only be accomplished through a limited number of embryo transfers, forcing all selection lines to go through a severe population bottleneck. A few generations after this transition and only for the lines that continued to be selected (DUK, DUC, DU6; see Table 1), family information and individual information was combined in a pedigree-based BLUP (Best Linear Unbiased Prediction)^12^ estimation of breeding values. Additionally, phenotyping in the line DU6 was massively extended by taking body weights at day 42 from all progeny, including females. After the implementation of these changes selection relied on estimated breeding values with a constraint on inbreeding.

### Sample collection and whole genome sequencing (WGS)

All animal procedures were performed in accordance with national and international guidelines and approved by the Animal Protection Board of the Institute for Farm Animal Biology. Genomic DNA was purified from tail biopsy samples using QIAamp DNA Mini Kit (Qiagen, Hilden, Germany) according to manufacturer’s recommendations. A total of 25 animals per line (150 animals in total) were sampled at two different time-points (Table 1). For the first time-point, females with the lowest kinship coefficient were chosen. Kinship was determined using the program INBREED implemented in the software SAS/STAT® (v9.4, SAS Institute Inc., USA). For the second time-point, females were chosen at random since the kinship coefficient is similar among subjects of the same line.

Library preparation and sequencing were carried out at the Competence Centre for Genomic Analysis (Kiel). Paired-end sequencing libraries were prepared using the TruSeq Nano DNA Library Prep kit following the manufacturer’s specifications (Illumina Inc., San Diego, CA, USA). Out of the 150 libraries, 60 were sequenced on a HiSeq 4000 platform (Illumina Inc.), and 90 samples were sequenced on a NovaSeq 6000 (Illumina Inc.) platform. The target coverage was 30× (high coverage set) and 5× (low coverage set), respectively. Read length was 151 nucleotides.

### Analysis of WGS data

Adapter removal and quality trimming were done using Trimmomatic v0.38^13^ for HisSeq reads and FASTP v0.19.6^14^ for NovaSeq reads. Read quality, was evaluated before and after processing with FastQC v0.11.5^15^. Reads were aligned to the mouse genome build GRCm38.p6^16^ from Ensembl version 93^17^ using the Burrow-Wheeler Aligner software in MEM mode (BWA-MEM)^18^ coupled with SAMtools v1.5^19^ in order to store alignments as BAM files. Per sample BAM files were processed sequentially with Picard tools^20^ by adding read group information (*AddOrReplaceReadGroups*), merging alignments from different read groups (*MergeSamFiles*), and by sorting (*SortSam*) and marking duplicated (*MarkDuplicates*) reads.

### Short variant calling and annotation

SNPs were detected according to GATK’s best practices for germline short variant discovery (GATK v 4.0.6.0)^21–24^. Systematic errors in base quality were corrected using *BaseRecalibrator* and dbSNP^25^ version 150 for *Mus musculus* (Ensembl version 93^17^). For each sample, variants were called with *HaplotypeCaller* and then combined with *GenomicsDBImport*. Joint genotyping was done with *GenotypeGVCFs* and then only bi-allelic variants (SNPs and INDELs) were retained. Filtering was applied separately for SNPs and INDELs. Site-level filtering was done following the Variant Quality Score Recalibration (VQSR) procedure. This comprised an internal variant set used as truth-training resource, created after stringent site-level filtering of the bi-allelic variants obtained from joint genotype calling, plus an external pre-filtered training variant set provided by the Mouse Genomes Project (MGP version 5^26^). Variants were genotyped as missing if the depth of coverage was either too low (<4), too high (3 standard deviations higher than the sample mean depth), or if the genotype quality (GQ) was too low (<20). The final set consisted of variants present in at least 15 samples per line (except for DU6 that had a lower coverage, so this threshold was lowered to 12 samples). Annotations were done using SnpEff v4.3t^27^ and missense mutations were further evaluated with Ensembl Variant Effect Predictor (VEP)^28^ to obtain their corresponding SIFT scores^29^ and to predict amino acid changes affecting protein function.

### Structural variant calling and annotation

Processed BAM files used for short variant calling were also used to detect large structural variants (SVs). Because of the considerable difference in coverage of the two sequence data sets, this was done independently for the high and the low coverage set. Three SV callers (Manta v.1.6.0^30^, Whamg v.1.7.0^31^ and Lumpy v.0.2.13^32^) were applied per line and per coverage set yielding six call sets per line.

Specific filters were applied depending upon the call set. SVs detected by Manta were site-filtered by excluding SVs with poor mapping quality (MAPQ < 30) or with excessive coverage (>3 × the median chromosome depth) that could be due to reads originated from low complexity regions. For each sample, only SVs with GQ ≥ 20 and read depth ≥5× were accepted. Whamg SV calls with sizes <50bp and >2Mb were filtered out to improve call accuracy. Here too, only calls with read depth ≥ 5× were accepted. Calls with GQ < 20 were filtered out. To reduce the number of false positive calls, high cross-chromosomal mappings were excluded, as Whamg is aware of but does not specifically call translocations. Likewise, SVs in poorly mapped regions were also removed. Lumpy SV calls for which supporting evidence (FORMAT/SU field) was below 5 (SU<5) were excluded, as well as SV calls with GQ<20. Since both Whamg and Lumpy do not have a built-in genotyping module, SV call sets were genotyped with Svtyper v0.7.1^33^ prior filtering for genotype quality. For each line and coverage set, SVs called by at least two SV callers were merged using Survivor v.1.0.7^34^ and kept if they were found in at least 10 samples. The final set consisted of the union of SVs detected in the high and low coverage read sets. We then intersected SV calls among all six mouse lines to obtain SVs private for each line (line-specific) and SVs shared among lines. SVs were annotated with VEP v.101.0^28^ focusing on variants affecting protein-coding genes with the maximum SV size set to 200 Mb. Functional classification was conducted after thorough literature and database search (OrthoDB v10^35^, Uniprot^36^, NCBI Entrez gene^37^), plus Gene Ontology enrichment analysis (Shiny GO^38^, false discovery rate [FDR] < 0.05). To further minimize false positives, SV calls overlapping gaps and high coverage regions (>80×) in the reference genome assembly were filtered out.

### Population genetics analysis

Genetic structure among all 150 samples was assessed using principal component analysis (PCA), hierarchical cluster (HC) analysis and genetic admixture. PCA and HC were computed using SNPRelate v1.22.0^39^. The ape v5.0 package was used for visualization of HC results^40^. Genetic admixture was estimated with ADMIXTURE v1.3.0^41^ after transforming the VCF file into a BED file using PLINK v2.00a2LM^42,43^. Linkage disequilibrium (LD) was evaluated after thinning the main VCF file with vcftools v0.1.13^44^ retaining sites at least 100Kb apart and then calculating r^2^ within windows of 5Mb using PLINK v2.00a2LM ^42,43^. Runs of homozygosity were estimated using the RoH extension^45^ in SAMtools/BCFtools v1.5^19^.

### Genetic differentiation and diversity analysis

The genomes of the trait-selected lines were compared to the neutrally evolving control line (FZTDU). For this, genetic differentiation was estimated using the F_ST_ index^46^ in sliding window mode (size=50Kb, step=25Kb, min 10 SNPs) using vcftools v0.1.13^44^. At each window, the arithmetic mean of the SNP-specific F_ST_ was calculated and then transformed into z-scores to represent its departure from the genomic mean. Additionally, all samples of the two fertility lines (DUK and DUC) were combined (pseudo-line: FERT) and compared to FZTDU as well. Since autosomes and the X-chromosomes have different effective population sizes, the X chromosome was standardized individually. In order to identify regions of distinct genetic differentiation (RDDs), F_ST_ windows appearing simultaneously in the 95^th^ percentile of a given contrast and in the bottom 10^th^ percentile of all other contrast were identified. These thresholds were chosen after testing multiple combinations of ≥95^th^ percentiles and ≤10^th^ percentiles, choosing the combination in which RDDs could be found in all contrasts. Genome-wide diversity patterns were assessed by measuring the nucleotide diversity (π)^47^ in sliding windows of 50Kb size (step size = 25Kb) using vcftools v0.1.13^44^.

### Gene annotation and enrichment analysis

Genes overlapping RDDs were identified using GenomicRanges^48^ and Ensembl 93’s^17^ *Mus musculus* gene set. In order to sort out the most relevant genes for each of the selected traits, thorough inspection of functional annotations, literature and SNP effects was conducted. This also included testing for enrichment of Gene Ontology Biological Processes (GOBP)^49,50^ and KEGG pathways^51–53^ using WebGestallt^54^. A FDR threshold of 10% was used as cutoff for significant enrichment of a term or pathway. Finally, genes in quantitative trait loci (QTLs) were identified by finding overlaps with QTL data compiled in the Mouse Genome Database^55^.

### Data handling, visualization and accessibility

Data processing and visualizations were done using R^56^ and the tidyverse package^57^. Raw sequencing data were deposited on the European Nucleotide Archive (accession: PRJEB44248). Scripts used to generate the results of this publication are available under https://github.com/sosfert/mmu_dummerstorf_wgs.

## Results and discussion

### Phenotypic impact of selection

Over the course of more than 140 generations, the selected traits have shown remarkable increments in each line (Fig. 1). The span and number of generations makes the present study the longest selection experiment ever reported in mice. Relative to the unselected control line FZTDU (exposed to genetic drift only), reproductive performance has doubled in DUK and DUC (Fig. 1A–B, F–G). Even though these two trait-selected lines have achieved comparable litter sizes at first delivery (>20 offspring)^58^, their reproductive lifespan differs, with 5.8 and 2.7 litters in average per lifetime for DUK and DUC, respectively^58^. A remarkable level of divergence has been achieved by the increased body size lines (Fig. 1C–D). DU6 individuals have almost tripled their weight compared to FZTDU (Fig. 1H), whereas mice of the protein line DU6P not only have become larger and heavier than FZTDU mice, but their level of muscularity is also considerably higher (Fig. 1D, 1I). In terms of running distance capacity, DUhLB mice can on average cover distances three times as long as those covered by FZTDU (Fig. 1J). With the exception of the obese line DU6^59^, each one of the trait-selected mouse lines has developed an extreme phenotype without obvious detrimental effects on their general health, well-being and longevity.

### WGS analysis and short variant detection

After quality filtering and trimming, >90% of the raw reads were mapped to the genome as pairs, with a mean insert size of ~380bp. For samples sequenced at a target coverage of 30×, mean genome-wide coverage averaged ~24×, with ~95% of genome territory covered at least 5×; samples sequenced at a target coverage of 5× averaged ~8× and ~72%, respectively (for a summary across all samples see Table 3).

**Table 3.**
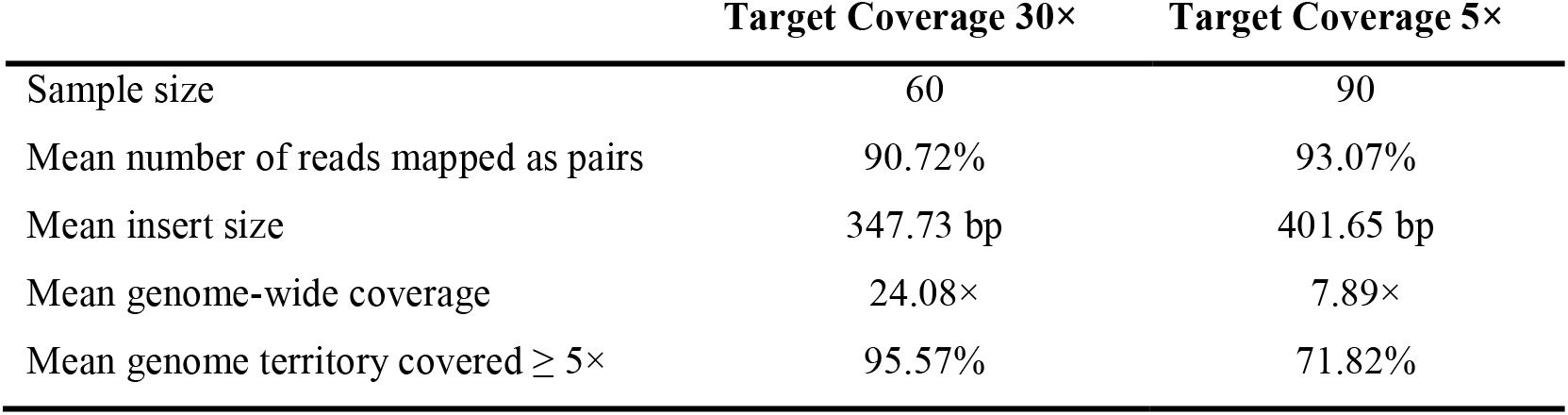
Summary metrics WGS data.

|  | Target Coverage 30× | Target Coverage 5× |
| --- | --- | --- |
| Sample size | 60 | 90 |
| Mean number of reads mapped as pairs | 90.72% | 93.07% |
| Mean insert size | 347.73 bp | 401.65 bp |
| Mean genome-wide coverage | 24.08× | 7.89× |
| Mean genome territory covered $\geq 5\times$ | 95.57% | 71.82% |

The final variant call set contained 5,099,945 SNPs and 896,078 INDELs (425,687 insertions; 470,391 deletions). The proportion of SNPs and INDELs overlapping dbSNP was 95% and 55%, respectively. This discrepancy is not necessarily due to a high number of artifacts in the INDEL set, but rather by the fact that INDELs are a much less characterized type of genetic variant in comparison^60^. Though considerably fewer variants were found in the trait-selected lines when compared with the control line FZTDU (Table 4), the number of SNPs was sufficient in all lines to conduct genome-wide selection scans (~ 1 SNP/Kb)^61^. Moreover, by retaining SNPs genotyped in at least 15 samples per line, the sample size required to have enough power to detect footprints of selection is sufficient as well^61^.

**Table 4.** Number of short variants discovered in each line.

| Line | SNPs | % in FZTDU | INDELs (Insertions + Deletions) | % in FZTDU |
| --- | --- | --- | --- | --- |
| DUK | 2,305,349 | 92.76 | 389,077 (186,193 + 202,884) | 89.46 |
| DUC | 2,615,584 | 92.76 | 431,296 (207,084 + 224,212) | 90.34 |
| DU6 | 2,744,788 | 93.18 | 453,449 (218,016 + 235,433) | 90.16 |
| DU6P | 2,899,902 | 91.04 | 479,457 (230,908 + 248,549) | 88.64 |
| DUhLB | 3,196,655 | 92.26 | 524,302 (252,965 + 271,337) | 89.84 |
| FZTDU | 4,453,865 | -- | 734,472 (356,672 + 377,800) | -- |

The number of alleles present in all six lines was ~1M, but very few alleles were shared by the trait-selected lines only (~3.3K) (Supplementary Fig. 1). The lines DU6P and DUhLB were the most polymorphic of the trait-selected lines, followed by the DU6. The two fertility lines (DUK, DUC) were the least polymorphic ones. Each of the trait-selected lines still shared >88% of its variants with the control line FZTDU (Table 4), indicating that despite genetic drift, the control line preserves most of the alleles that were present when the selective breeding process started and that it is a reliable proxy of the original founder population.

Almost all SNPs and INDELs (~97%) occurred in non-coding regions (introns: ~56%; intergenic: ~41%). This is not an unexpected outcome considering that only ~2% of the genome codes for proteins and genetic variation is wide-spread. Inter-genic variants could affect regulatory elements of gene expression, as well as transcripts not yet described^62^, whereas intronic variants could affect gene splicing^63^.

Based on assessment of variant annotations, a very small number of variants (20,236 SNPs and 2,387 INDELs) were classified as high-impact and moderate-impact mutations, and could interfere with gene transcription or translation. These “impact-variants” were screened for i) being private for any trait-selected line (Supplementary Table 1) and ii) the functional categories their affected genes belonged to. For the lines DUC, DU6 and DU6P there were significantly enriched functions that are coherent with the selected traits, including metabolic pathways (DU6), anabolism and regulation of protein synthesis (DU6P) and embryonic development (DUC). For both DUK and DUhLB, results were less explicit, with most GO terms relating to immunity (Supplementary Data 1).

### Runs of homozygosity and linkage disequilibrium

While for the five trait-selected lines, most of the SNP loci (57.5% - 81.5%) were already fixed for either the reference or the alternative allele, in the control line FZTDU alleles were mostly (>75%) polymorphic (Supplementary Fig. 2). This disparity was also reflected by the distribution of frequencies for the alternative allele, displaying a “U” shape that is much more pronounced in the trait-selected lines than in the control line (Supplementary Fig. 3). Genomes of mice from the control line FZTDU also had higher nucleotide diversity (Supplementary Fig. 4 and 5). Accordingly, runs of homozygosity (RoH) covered between ~65% and ~78% (~50% as 1-8Mb tracts) of the genome length of the trait-selected lines, but only ~45% (~23% as 1-8Mb tracts) of the genome length of FZTDU (Fig. 2A). Analysing RoH shared among individuals of a population can aid to detect past selection events^64^; however, this is applicable as long as RoH events are rare in the genome (RoH islands), which is not the case here, where RoH are widespread, indicating that the observed degree of homozygosity is the result of a combination of multiple evolutionary forces.

**Fig. 2.**
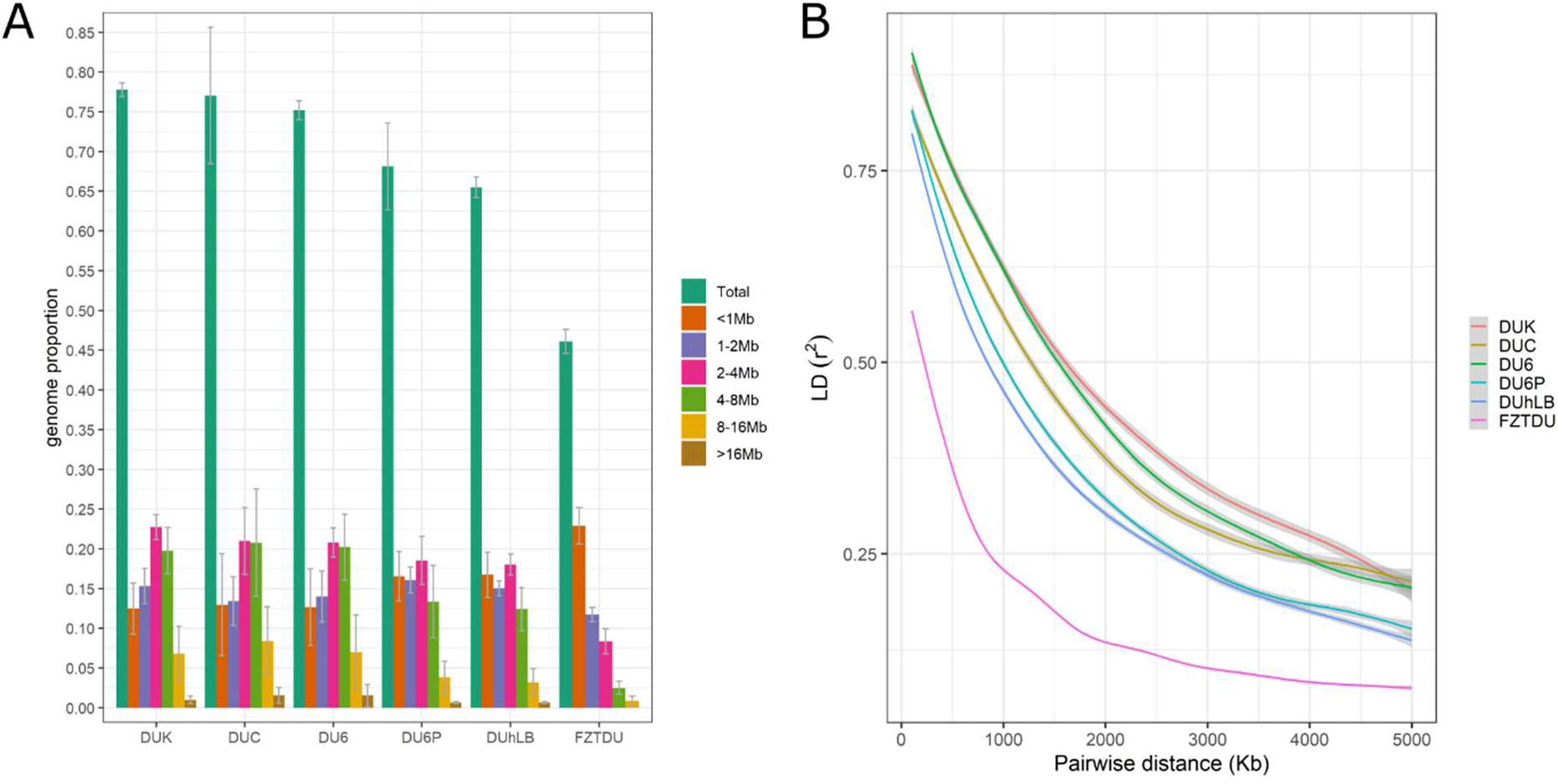
Runs of homozygosity (RoH) and linkage disequilibrium decay in the Dummerstorf mouse lines. (**A**) Per line average extent of homozygosity as a fraction of the genome length. RoH of different length range are specified by colours. Error bars show ±1SD. (**B**) Decay of the mean genotype correlations among SNP pairs as close as 0.1Mb and as far as 5Mb.

Linkage disequilibrium decay can be classified into three decaying patterns with decreasing decay strength; one for the three most homozygous trait-selected lines (DUK, DUC and DU6; upper three lines Fig. 2B), a second for the two least homozygous trait-selected lines (DU6P and DUhLB; middle two lines Fig. 2B) and a third for the unselected line FZTDU (bottom line Fig. 2B). Overall, linkage disequilibrium decay clearly differs between trait-selected lines and FZTDU. Such extensive linkage disequilibrium has been previously reported in mountain Gorillas, in which the level of inbreeding due to population decline has led to similar linkage disequilibrium patterns as the ones found in the trait-selected lines^65^.

### Population structure of the Dummerstorf mouse lines

The genetic relationship among the 150 Dummerstorf mice was assessed by PCA, HC and admixture analysis using the 5,099,945 SNPs obtained by variant calling. Samples formed a hierarchical group structure that represented each of the Dummerstorf lines and also distinguished trait-selected from unselected control animals (Fig. 3A–C). There was no admixture present in the trait-selected lines, except for one DUC animal sharing ancestry with mice from DU6P (Fig. 3D). FZTDU is represented as an admixture of all the trait-selected lines with similarly large contributions of the four older lines and a significantly larger contribution of DUhLB. This is expected because this mouse line is the youngest and has had the least number of generations that underwent selection (Fig. 3D).

**Fig. 3.**
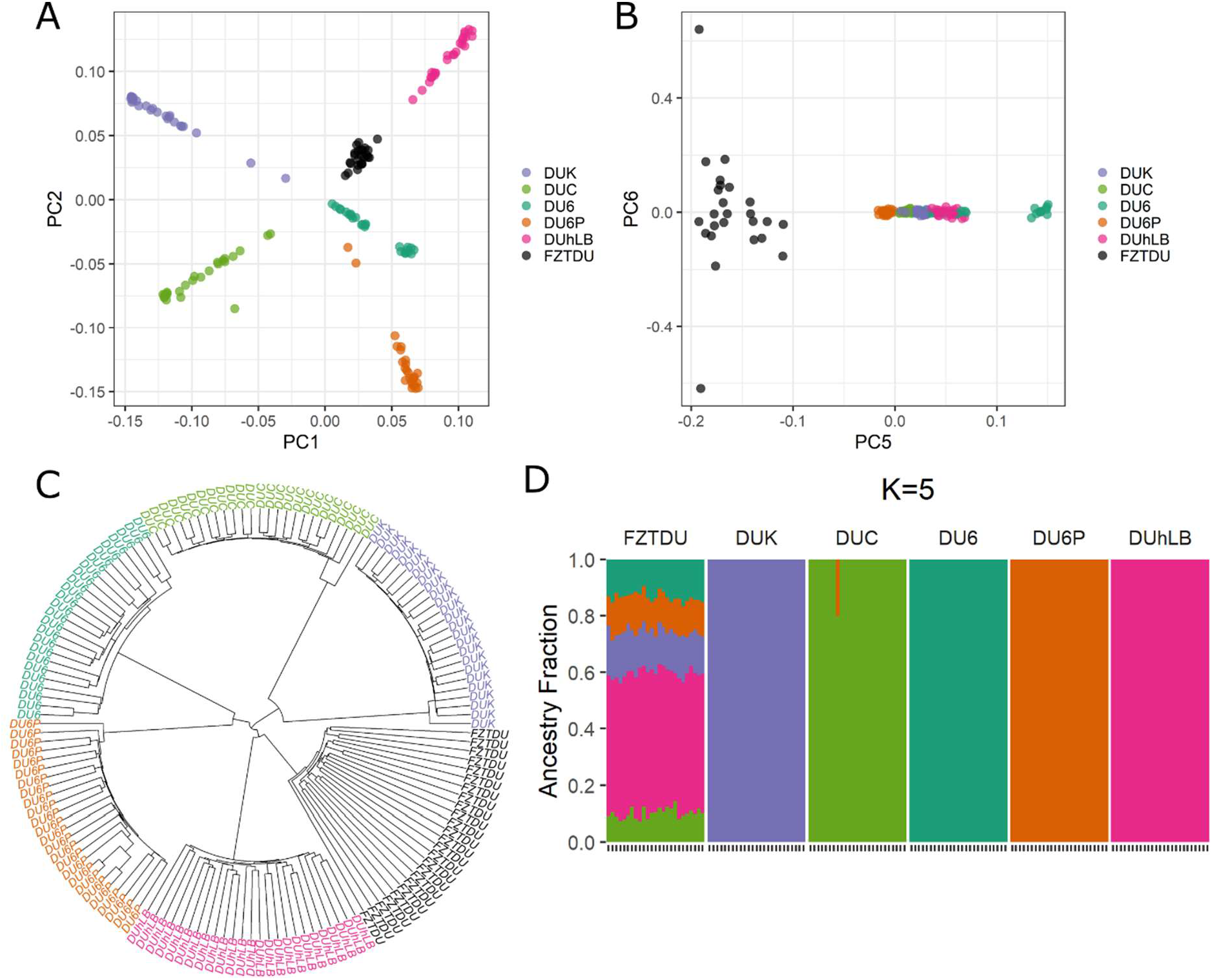
Genetic structure and cluster assignment of 150 mice of the six Dummerstorf mouse lines. (**A**) Scores for principal components 1 and 2 separating individuals into distinctive mouse line clusters. (**B**) Scores for principal components 5 and 6 separating the trait-selected lines from the control line FZTDU. (**C**) Hierarchical clustering analysis. (**D**) Genetic composition of each mice (indicated by 25 ticks on the x-axis) in terms of the five trait-selected lines. Individuals are coloured according to the respective line of origin.

### Genetic differentiation of the trait-selected lines

Mean genome-wide pairwise genetic differentiation among trait-selected lines estimated by F_ST_ ranged from 0.44 to 0.61 (Fig. 4B).The highest level of differentiation was found between either one of the fertility lines and the body mass line DU6 (F_ST(DUK-DU6)_ = 0.61 and F_ST(DUC-DU6)_ = 0.59; Fig. 4B), followed by the differentiation between the two fertility lines themselves (F_ST(DUK-DUC)_ = 0.57; Fig. 4B). Although pairwise genetic differentiation between trait-selected lines and the control line was similar in all comparisons (F_ST_ ~ 0.3), it was lowest in the pairwise comparison between the two most polymorphic lines (F_ST(DUhLB-FZTDU)_ = 0.26; Fig. 4B). Such strong levels of differentiation occur mainly as a result of reproductive isolation and genetic drift^66^; however, it is expected that a subset of alleles that have arrived to fixation due to selection contribute to genetic differentiation as well. The challenge is thus to sort out which genomic regions contain such beneficial alleles.

**Fig. 4.**
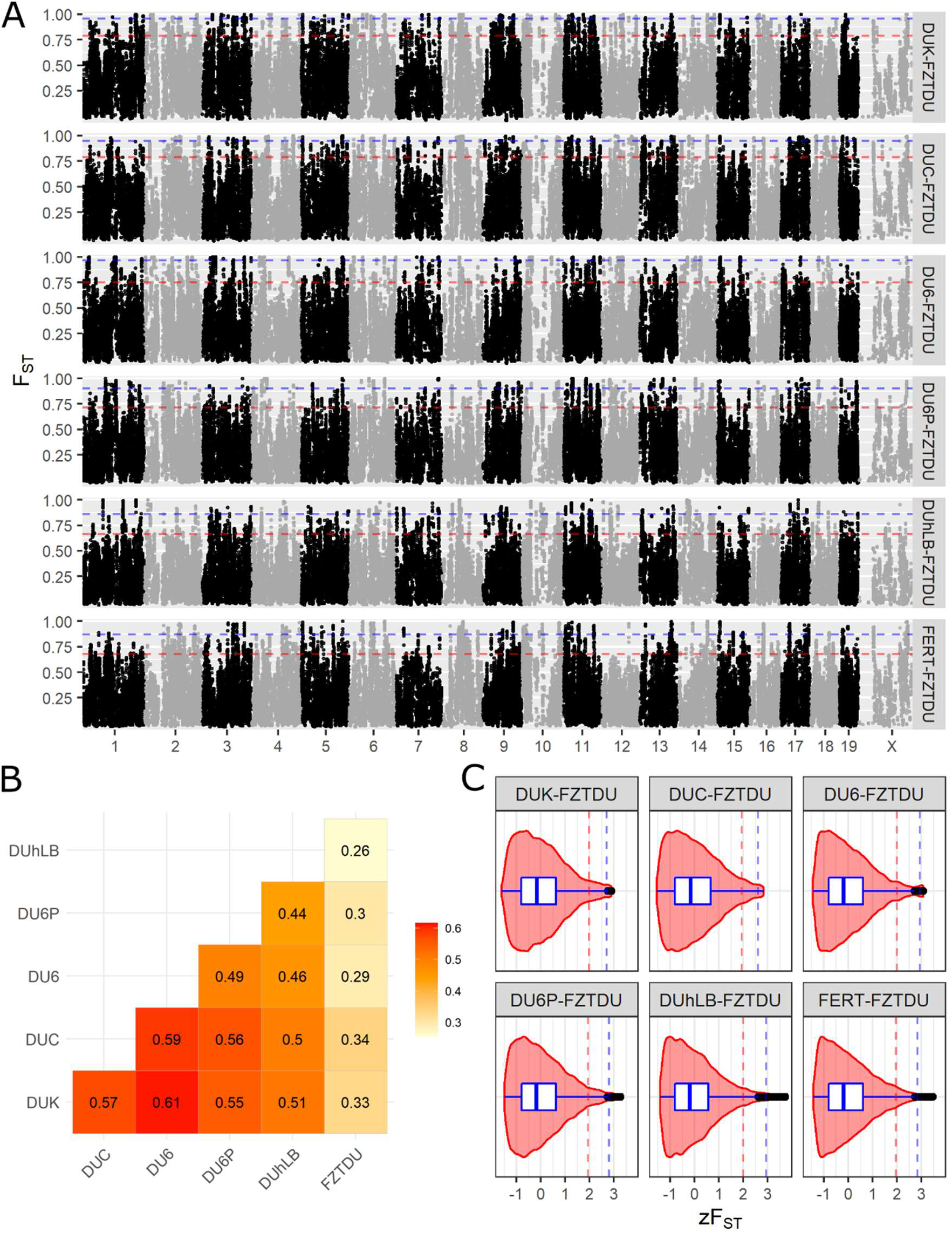
Genetic differentiation of the Dummerstorf trait-selected lines. (**A**) Genome-wide scans of genetic differentiation in sliding-window mode (size=50Kb, step=25Kb) contrasting each trait-selected line to FZTDU. Each window is the average F_ST_ of at least 10 SNPs. (**B**) Pairwise genomic mean F_ST_ among all six Dummerstorf lines. (**C**) F_ST_ distribution as z-scores, illustrating the departure of each window from the mean genomic level of genetic differentiation. Dotted lines indicate the 95^th^ (red) and 99^th^ (blue) percentiles.

### Trait-specific regions of genetic differentiation

Genome-wide scans were conducted in order to detect genomic regions of consistent genetic differentiation between each trait-selected line and FZTDU. The pseudo-line FERT was also included, for a total of six F_ST_ contrasts. Choosing genomic regions of interest by focusing on the most differentiated regions (percentile 95^th^ or 99^th^ of the F_ST_ distribution) resulted in the detection of multiple loci in every chromosome (Fig. 4A). Because these regions were frequent and did not sufficiently depart from the global level of differentiation to be considered genomic outliers (i.e. max. zF_ST_: 2.89 - 3.47), a more stringent approach was applied to identify line-specific regions of high genetic differentiation, while reducing the influence of genetic drift. These regions of distinct genetic differentiation (hereafter referred to as RDDs) appeared simultaneously in the top 5% F_ST_ windows of the target contrast and in the bottom 10% of all the remaining contrasts, occurring close to each other in only a subset of chromosomes (Fig. 5A–C, Fig. 6A–C) and containing multiple genes (Supplementary Data 2-7), some of which were related to the selected traits (see below).

**Fig. 5.**
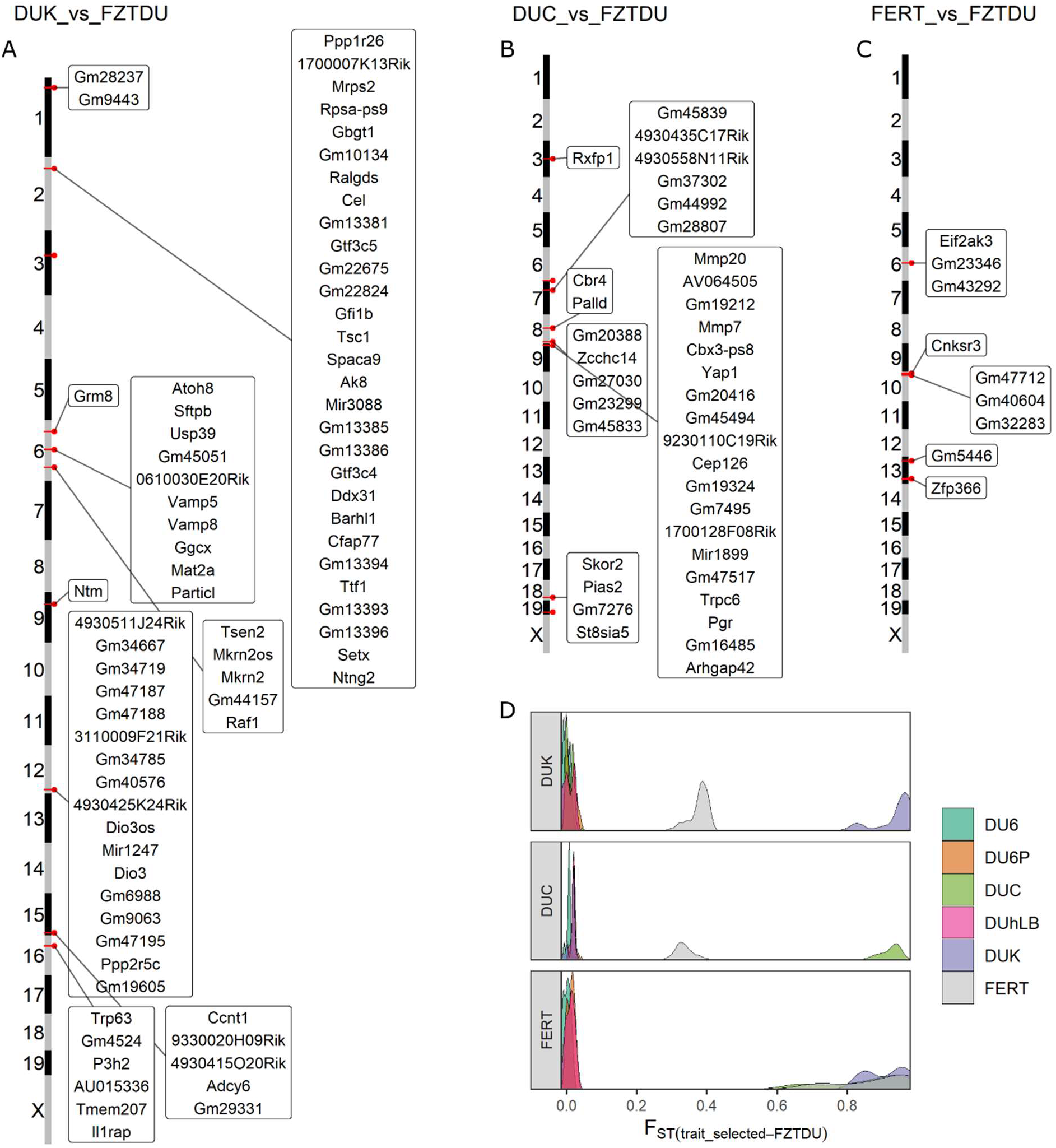
Genes mapped to regions of distinct genetic differentiation (RDD) for fertility lines. (**A**-**C**) Genomic overview of RDDs for each of the fertility lines (DUK, DUC) and the joint pseudo-line (FERT). (**D**) F_ST_ distribution of RDDs, demonstrating the gap in F_ST_ between the target lines and the rest.

**Fig. 6.**
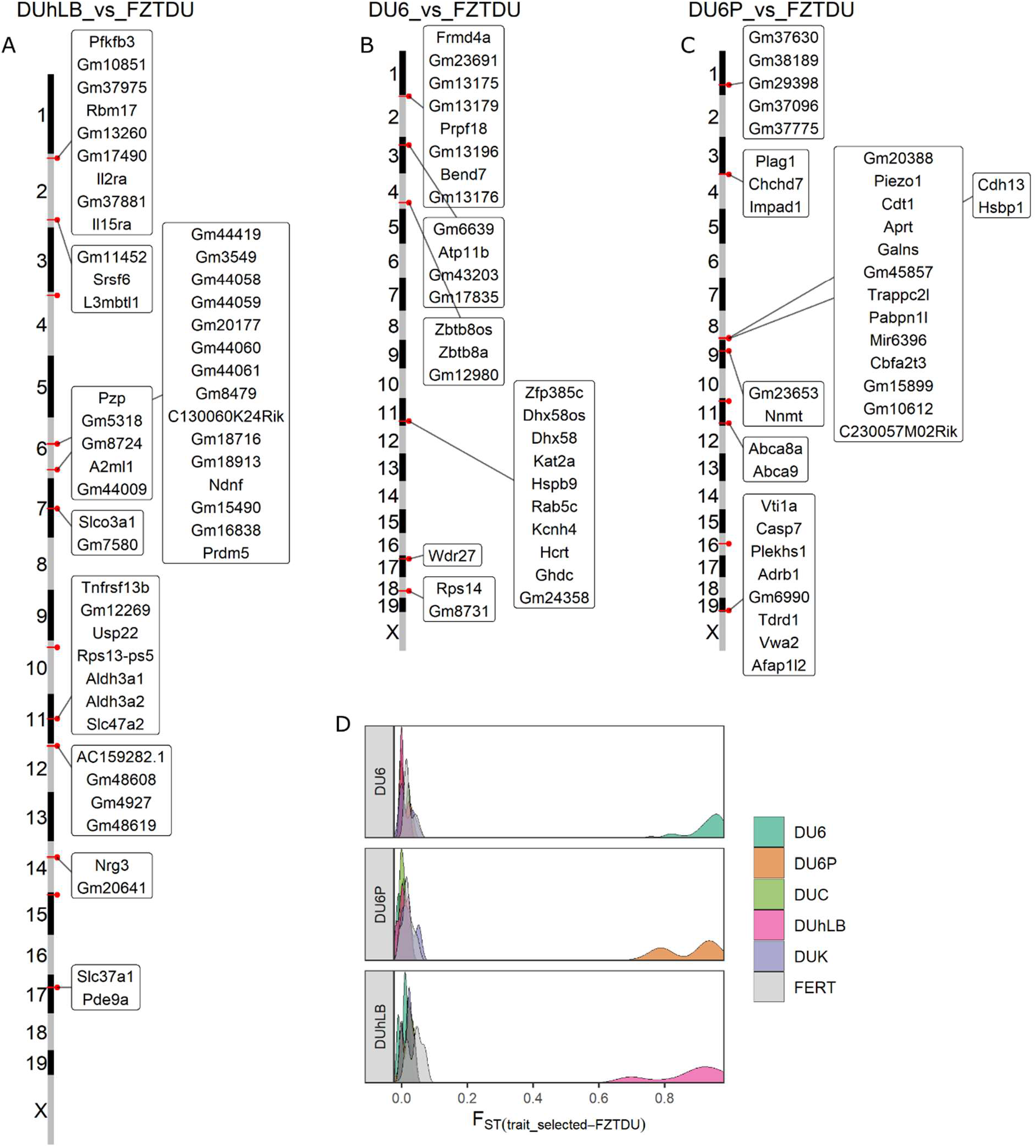
Genes mapped to regions of distinct genetic differentiation for the body mass lines (DU6 and DU6P) and the treadmill running endurance line (DUhLB). (**A**-**C**) Genomic overview of RDDs for DUhLB, DU6 and DU6P, respectively. (**D**) F_ST_ distribution of RDDs, demonstrating the gap in F_ST_ between the target lines and the rest.

### Line-specific patterns of structural variation

Despite primarily thought to be deleterious and implicated in disease phenotypes^67^, large chromosomal rearrangements such as deletions, duplications and inversions have an important role in local adaptation and divergence of populations^68^. These structural variants can lead to gene expression differences by disrupting genes and altering gene dosage^69^. Because copy number variation often results in notable phenotypic differences it is likely a subject to selection during domestication^70^. For example, genes related to metabolic activity and production traits have been shown to be affected by copy number variation during artificial selection of cattle^71^, goats^72^ and pigs^73^.

After calling and filtering, only duplications, deletions and inversions remained in the final SV data set. Insertions did not occur in enough samples to be included in the analysis. SVs were predominantly located in non-coding regions (98%) where they could affect gene expression. Also, SVs were more abundant in the trait-selected lines (5,195 - 6,856 SVs) than in the control line (4,521 SVs) implying that large genomic rearrangements could contribute to the development of the selected traits. In order to associate SVs to each selected trait, line-specific SVs overlapping protein coding genes were identified and characterized in greater detail (Supplementary Data 8). The total number of these line-specific SVs ranged from 9 (FZTDU) to 36 (DUC), comprising mostly deletions and inversions (Table 5). Most SVs were polymorphic and large length differences were observed between polymorphic and fixed SVs (Supplementary Table 3). Fixed line-specific deletions were detected in all lines, whereas duplications were found only in DU6P, and inversions only in DUC, DU6P and DUhLB (Supplementary Table 4).

**Table 5.** Summary of structural variants detected in all mice lines.

|  | Total |  |  |  | Line-specific-genic |  |  |  |
| --- | --- | --- | --- | --- | --- | --- | --- | --- |
|  | DEL | DUP | INV | Total | DEL | DUP | INV | Total |
| DUK | 4,633 | 32 | 530 | 5,195 | 11 | 2 | 7 | 20 |
| DUC | 5,560 | 48 | 1,248 | 6,856 | 10 | 2 | 24 | 36 |
| DU6 | 5,025 | 27 | 551 | 5,603 | 11 | 0 | 7 | 18 |
| DU6P | 4,339 | 23 | 2,091 | 6,453 | 9 | 1 | 14 | 24 |
| DUhLB | 4,614 | 20 | 1,508 | 6,142 | 10 | 1 | 9 | 20 |
| FZTDU | 3,902 | 14 | 605 | 4,521 | 4 | 0 | 5 | 9 |

The number of genes affected by fixed line-specific SVs varied from 1 (DUC, DU6P, FZTDU) to 5 (DUK), but went up to more than a thousand for genes affected by large polymorphic inversions (Supplementary Table 5). These genes were classified in functional groups based on the biological processes they are associated with (Supplementary Table 6). The most gene-rich functional groups are the ones associated with sensory perception, predominantly olfaction (found in the fertility lines DUK and DUC), followed by “cell cycle and nucleic acid transcription and translation” (in DUC), and “metabolism and energy conversion” (DUC, DU6P).

### Genes associated with fertility

Genes detected in RDDs for DUK were enriched for “phospholipase D signalling pathway” (Supplementary Table 3). In granulomas cells, phospholipase D activity is stimulated by GnRH, thereby inducing or inhibiting cell differentiation depending on the maturation state of the ovarian follicle^74^. Other genes encode for proteins involved in the ovarian development and maintenance of the primordial follicle reserve (*Tsc1*^75^, *Trp63*^76^), in the vascularization of the placenta (*Atoh8*^77^) and facilitate maternal supplied lipids and dietary fat digestion in neonatal mice (*Cel*^78,79^). Furthermore, DUK shares a fecundity associated region (*Sftpb*, *Usp39*, *Tmem150*, *Rnf181*, *Vamp5*, *Vamp8*, *Cgcx*, *Mat2a*) with Qsi5 mice^80^, an inbred mouse line known for its increased litter size, and candidate genes associated with birth rate and male fertility in humans (*Ntm*^81^) and litter size in cattle, goats and pigs (*Dio3*^82–84^). Interestingly, analysis of private SVs detected a 317bp deletion affecting *Olfr279* (Supplementary Data 8). This gene has been associated to mouse male sub-fertility^85^ and more generally, olfactory receptors could regulate fertilization^86,87^.

Significantly enriched terms for DUC included “intracellular steroid hormone receptor signalling pathway”, involving progesterone receptor (*Pgr*), an established reproductive gene, which carries a missense mutation that is fixed in and specific for DUC (Supplementary Fig. 6). In line with the increased progesterone levels in DUC females^88^, the progesterone receptor gene in DUC harbors a missense SNP mutation with high genetic differentiation (F_ST_ ~ 0.93). The amino acid substitution is predicted to compromise the protein’s biochemical properties (SIFT score = 0.04). Interestingly, a Neanderthal missense mutation in *Pgr* associated with increased fertility was recently reported to segregate in human populations^89^. Further candidates in DUC control ovarian follicle development, uterine growth, and endometrial angiogenesis during pregnancy (*Yap1*^90^, *Rxfp1*^91,92^). In the context of preparation of the endometrium for implantation and pregnancy and progesterone signalling, the gene *Rrm2*^92^ was identified by the structural variation of the DUC genome.

The fertility lines DUK and DUC have been bred according to the same criteria, share the same evolutionary history, and both have been able to more than double the number of pups per litter since the beginning of selection. Despite these commonalities, improved fertility is achieved via different physiological pathways in each line^88^. For example, females from both fertility lines have an increased ovulation rate, but only DUK exhibits follicles containing multiple oocytes; DUC on the other hand shows an increased progesterone level compared to DUK and FZTDU^88^. The scarce number of RDDs in the combined FERT population also illustrates this discrepancy. Candidate RDD and line-specific SV overlapping genes in both fertility lines likely affect the reproductive process on multiple levels such as ovarian physiology, placentation, sex steroid signaling and milk composition.

### Genes associated with body size and endurance

Two of the Dummerstorf trait-selected mouse lines have increased their body weight in response to selection. The “giant” DU6 line (selected for body mass at 6 weeks of age) exhibits an obese phenotype^7^ while the protein-mass line DU6P (selected for protein mass in the carcass) is lean and muscular^93^. In line with the obese phenotype, DU6 candidate genes overlapping RDDs regulate energy metabolism and food intake (*Hcrt*^94^) and are linked to feed efficiency and body composition in other species (*Atp11*^95^, *Wdr27*^96^). On the other hand, DU6 mice also exhibit larger bones^59^ and the analysis of SVs detected *Smad5*, a modulator of bone formation^97^, to be partially overlapped by a heterozygous deletion and a heterozygous inversion. Candidate genes in the RDDs for DU6P conform with growth-related major quantitative trait loci found in sheep and are known to influence stature and body size in cattle, pigs and human (*Plag1*^98,99^, *Chchd7*^98–100^, *Impad1*^101^). In line with this, an SV (deletion) was found overlapping *Fam92a*, a gene that is involved in limb development^102^. Further candidates for lean body mass are the RDD overlapping genes *Piezo1* (myotube formation^103,104^) and *Cdh13* (control of lipid content in developing adipocytes^105–107^).

Finally, genes specific for the endurance line DUhLB participate in lipid metabolism (these animals manifest faster mobilization of lipids during exercise). Only two DUhLB genes (*Aldh3a1* and *Aldh3a2*, the later containing 3 missense SNPs (Supplementary Fig. 7C)) caused the significant enrichment of the “Histidine metabolism” and “beta-Alanine metabolism” pathways. The “marathon mice” DUhLB have developed a striking metabolic phenotype characterized by accelerated browning of subcutaneous fat and altered mitochondrial biogenesis in response to selection for high treadmill performance^108^. Likewise, detected RDD candidate genes are involved in the development of brown adipocytes (*Srsf6*^109^), removal of toxic waste products from lipid metabolism (*Aldh3a2*^110^), fatty acids mobilization, mitochondria content, and cristae complexity (*Il15r*^111^) and in the regulation of glycolysis associated to obesity and weight gain (*Pfkfb3*^112,113^). Moreover, SV analysis detected a ~2.8Kb inversion in *Atp5j* whose overexpression has been shown to counteract exercise-induced cardiac hypertrophy in mice^114^.

## Conclusions

The Dummerstorf trait-selected mouse lines have evolved in isolation through selective breeding and genetic drift, resulting in extensive long runs of homozygosity and ubiquitous regions of high genetic differentiation. Distinguishing between both evolutionary forces is a challenging task, which will require further research, but by focusing on regions of distinct genetic differentiation with respect to a control line that represents the founder population of this selection experiment, we were able to identify genes with important functions associated to the selected traits

Over the span of this selection experiment, traits have improved continuously and have not decayed despite the dramatic loss of genetic diversity within lines. This implies that many of the alleles that contribute to trait improvement have arrived to fixation and that these lines are highly enriched for such alleles. Therefore, a deeper understanding of the genomes of the trait-selected Dummerstorf mouse lines will provide valuable insights into the genetic basis of important polygenic traits and constitutes an unprecedented scientific resource for geneticists, physiologists and the wider biomedical research community.

## Acknowledgements

This study was funded by the Leibniz Collaborative Excellence programme (K52/2017, SOS-FERT). We acknowledge all funders and researchers who have made it possible for this experiment to continue over more than 50 years and to survive fundamental political changes, institute closures, reorganizations, and scientific reorientations. We particularly thank the staff of the FBN Service Group Lab Animal Facility for excellent animal care and cooperation. This work was supported by the DFG Research Infrastructure NGS_CC (project 407495230) as part of the Next Generation Sequencing Competence Network (project 423957469).

## Author contributions

SEPV conducted bioinformatic and population genomic analysis, interpreted the data and wrote the manuscript. HR provided genomic DNA material, interpreted the data and advised population genomic analysis. ML supervised mouse breeding, sample collection and phenotypic records. NR conceived the study, interpreted the data and supervised population genomic analysis and edited the manuscript. LD conducted structural variation analysis and wrote the manuscript accordingly. JF conceived the study, interpreted the data, supervised population genomic analysis and edited the manuscript. SQ advised population genomic analysis and provided suggestions to improve the manuscript. JW interpreted the data and edited the manuscript. SF conducted whole genome sequencing. GHS conducted whole genome sequencing and advised bioinformatics analysis. JS conceived the study, interpreted the data and wrote the manuscript. All authors have reviewed and approved the manuscript.

## Competing interests

The authors declare no competing interests.

## Notes

### Competing Interest Statement

The authors have declared no competing interest.

## References

1. Conner, J. K. Artificial Selection. in Encyclopedia of Evolutionary Biology (ed. Kliman, R. M. B. T.-E. of E. B.) 107–113 (Academic Press, 2016). doi:https://doi.org/10.1016/B978-0-12-800049-6.00053-6.

2. Kukekova, A. V. et al. Red fox genome assembly identifies genomic regions associated with tame and aggressive behaviours. Nat. Ecol. Evol. 2, 1479–1491 (2018).

3. Castro, J. P. et al. An integrative genomic analysis of the Longshanks selection experiment for longer limbs in mice. Elife 8, (2019).

4. Schueler, L. Mouse strain Fzt:DU and its use as model in animal breeding research. Arch. für Tierzucht (Archives Anim. Breeding) 28, 357–363 (1985).

5. Dietl, G., Langhammer, M. & Renne, U. Model simulations for genetic random drift in the outbred strain Fzt: DU. Arch. FUR TIERZUCHT 47, 595–604 (2004).

6. Langhammer, M. et al. Reproductive performance primarily depends on the female genotype in a two-factorial breeding experiment using high-fertility mouse lines. Reproduction 153, 361–368 (2017).

7. Renne, U. et al. Lifelong Obesity in a Polygenic Mouse Model Prevents Age- and Diet-Induced Glucose Intolerance– Obesity Is No Road to Late-Onset Diabetes in Mice. PLoS One 8, e79788 (2013).

8. Ohde, D. et al. Advanced Running Performance by Genetic Predisposition in Male Dummerstorf Marathon Mice (DUhTP) Reveals Higher Sterol Regulatory Element-Binding Protein (SREBP) Related mRNA Expression in the Liver and Higher Serum Levels of Progesterone. PLoS One 11, e0146748 (2016).

9. Justice, M. J. & Dhillon, P. Using the mouse to model human disease: Increasing validity and reproducibility. DMM Disease Models and Mechanisms vol. 9 101–103 (2016).

10. Rosenthal, N. & Brown, S. The mouse ascending: Perspectives for human-disease models. Nature Cell Biology vol. 9 993–999 (2007).

11. Walsh, B. & Lynch, M. Evolution and Selection of Quantitative Traits. vol. 1 (Oxford University Press, 2018).

12. Henderson, C. R. Best Linear Unbiased Estimation and Prediction under a Selection Model. Biometrics 31, 423–447 (1975).

13. Bolger, A. M., Lohse, M. & Usadel, B. Trimmomatic: a flexible trimmer for Illumina sequence data. Bioinformatics 30, 2114–2120 (2014).

14. Chen, S., Zhou, Y., Chen, Y. & Gu, J. fastp: an ultra-fast all-in-one FASTQ preprocessor. Bioinformatics 34, i884–i890 (2018).

15. Andrews, S. FastQC A Quality Control tool for High Throughput Sequence Data. http://www.bioinformatics.babraham.ac.uk/projects/fastqc/.

16. Waterston, R. H. et al. Initial sequencing and comparative analysis of the mouse genome. Nature 420, 520–562 (2002).

17. Yates, A. D. et al. Ensembl 2020. Nucleic Acids Res. 48, D682–D688 (2020).

18. Li, H. & Durbin, R. Fast and accurate short read alignment with Burrows-Wheeler transform. Bioinformatics 25, 1754–1760 (2009).

19. Li, H. et al. The sequence alignment/map format and SAMtools. Bioinformatics 25, 2078–2079 (2009).

20. Broad Institute. Picard Tools. http://broadinstitute.github.io/picard/.

21. McKenna, A. et al. The Genome Analysis Toolkit: a MapReduce framework for analyzing next-generation DNA sequencing data. Genome Res 20, 1297–1303 (2010).

22. Depristo, M. A. et al. A framework for variation discovery and genotyping using next-generation DNA sequencing data. Nat. Genet. 43, 491–501 (2011).

23. Auwera, G. A. et al. From FastQ data to high confidence variant calls: the Genome Analysis Toolkit best practices pipeline. Curr Protoc Bioinforma. 11, (2013).

24. Poplin, R. et al. Scaling accurate genetic variant discovery to tens of thousands of samples. bioRxiv 201178 (2017) doi:10.1101/201178.

25. Sherry, S. T. dbSNP: the NCBI database of genetic variation. Nucleic Acids Res. 29, 308–311 (2001).

26. Keane, T. M. et al. Mouse genomic variation and its effect on phenotypes and gene regulation. Nature 477, 289–294 (2011).

27. Cingolani, P. et al. A program for annotating and predicting the effects of single nucleotide polymorphisms, SnpEff: SNPs in the genome of Drosophila melanogaster strain w1118; iso-2; iso-3. Fly 6, 80–92 (2012).

28. McLaren, W. et al. The Ensembl Variant Effect Predictor. Genome Biol. 17, 122 (2016).

29. Ng, P. C. & Henikoff, S. SIFT: Predicting amino acid changes that affect protein function. Nucleic Acids Res. 31, 3812–3814 (2003).

30. Chen, X. et al. Manta: Rapid detection of structural variants and indels for germline and cancer sequencing applications. Bioinformatics 32, 1220–1222 (2016).

31. Kronenberg, Z. N. et al. Wham: Identifying Structural Variants of Biological Consequence. PLOS Comput. Biol. 11, e1004572 (2015).

32. Layer, R. M., Chiang, C., Quinlan, A. R. & Hall, I. M. LUMPY: A probabilistic framework for structural variant discovery. Genome Biol. 15, R84 (2014).

33. Chiang, C. et al. SpeedSeq: Ultra-fast personal genome analysis and interpretation. Nat. Methods 12, 966–968 (2015).

34. Jeffares, D. C. et al. Transient structural variations have strong effects on quantitative traits and reproductive isolation in fission yeast. Nat. Commun. 8, 1–11 (2017).

35. Kriventseva, E. V. et al. OrthoDB v10: Sampling the diversity of animal, plant, fungal, protist, bacterial and viral genomes for evolutionary and functional annotations of orthologs. Nucleic Acids Res. 47, D807–D811 (2019).

36. Bateman, A. et al. UniProt: The universal protein knowledgebase. Nucleic Acids Res. 45, D158–D169 (2017).

37. Maglott, D., Ostell, J., Pruitt, K. D. & Tatusova, T. Entrez gene: Gene-centered information at NCBI. Nucleic Acids Res. 39, D52–D57 (2011).

38. Ge, S. X., Jung, D., Jung, D. & Yao, R. ShinyGO: A graphical gene-set enrichment tool for animals and plants. Bioinformatics 36, 2628–2629 (2020).

39. Zheng, X. et al. A high-performance computing toolset for relatedness and principal component analysis of SNP data. Bioinformatics 28, 3326–8 (2012).

40. Paradis, E. & Schliep, K. ape 5.0: an environment for modern phylogenetics and evolutionary analyses in R. Bioinformatics 35, 526–528 (2018).

41. Alexander, D. H., Novembre, J. & Lange, K. Fast model-based estimation of ancestry in unrelated individuals. Genome Res. 19, 1655–1664 (2009).

42. Chang, C. C. et al. Second-generation PLINK: rising to the challenge of larger and richer datasets. Gigascience 4, 7 (2015).

43. Purcell, S. & Chang, C. PLINK 2. https://www.cog-genomics.org/plink/2.0/.

44. Danecek, P. et al. The variant call format and VCFtools. Bioinformatics 27, 2156–2158 (2011).

45. Narasimhan, V. et al. BCFtools/RoH: A hidden Markov model approach for detecting autozygosity from next-generation sequencing data. Bioinformatics 32, 1749–1751 (2016).

46. Weir, B. S. & Cockerham, C. C. Estimating F-Statistics for the Analysis of Population Structure. Evolution (N. Y). 38, 1358 (1984).

47. Nei, M. & Li, W. H. Mathematical model for studying genetic variation in terms of restriction endonucleases. Proc. Natl. Acad. Sci. U. S. A. 76, 5269–5273 (1979).

48. Lawrence, M. et al. Software for Computing and Annotating Genomic Ranges. PLoS Comput. Biol. 9, e1003118 (2013).

49. Ashburner, M. et al. Gene ontology: tool for the unification of biology. The Gene Ontology Consortium. Nat. Genet. 25, 25–29 (2000).

50. The Gene Ontology Resource: 20 years and still GOing strong. Nucleic Acids Res. 47, D330–D338 (2019).

51. Kanehisa, M. & Goto, S. KEGG: kyoto encyclopedia of genes and genomes. Nucleic Acids Res. 28, 27–30 (2000).

52. Kanehisa, M., Sato, Y., Furumichi, M., Morishima, K. & Tanabe, M. New approach for understanding genome variations in KEGG. Nucleic Acids Res. 47, D590–D595 (2019).

53. Kanehisa, M. Toward understanding the origin and evolution of cellular organisms. Protein Sci. 28, 1947–1951 (2019).

54. Liao, Y., Wang, J., Jaehnig, E. J., Shi, Z. & Zhang, B. WebGestalt 2019: gene set analysis toolkit with revamped UIs and APIs. Nucleic Acids Res. 47, W199–W205 (2019).

55. Bult, C. J. et al. Mouse Genome Database (MGD) 2019. Nucleic Acids Res. 47, D801–D806 (2019).

56. R Core Team. R: A Language and Environment for Statistical Computing. (R Foundation for Statistica Computing, 2020).

57. Wickham, H. et al. Welcome to the {tidyverse}. J. Open Source Softw. 4, 1686 (2019).

58. Langhammer, M. et al. Two mouse lines selected for large litter size display different lifetime fecundities. Reproduction (2021) doi:10.1530/REP-20-0563.

59. de-Diego, I. et al. Titan mice are unique short-lived mammalian model of metabolic syndrome and aging. bioRxiv 2020.05.11.088625 (2020) doi:10.1101/2020.05.11.088625.

60. Hu, J. & Ng, P. C. Predicting the effects of frameshifting indels. Genome Biol. 13, R9 (2012).

61. Ma, Y. et al. Properties of different selection signature statistics and a new strategy for combining them. Heredity (Edinb). 115, 426–436 (2015).

62. Bartonicek, N. et al. Intergenic disease-associated regions are abundant in novel transcripts. Genome Biol. 18, 241 (2017).

63. Scotti, M. M. & Swanson, M. S. RNA mis-splicing in disease. Nature Reviews Genetics vol. 17 19–32 (2016).

64. Kim, E.-S. et al. Effect of Artificial Selection on Runs of Homozygosity in U.S. Holstein Cattle. PLoS One 8, e80813 (2013).

65. Xue, Y. et al. Mountain gorilla genomes reveal the impact of long-term population decline and inbreeding. Science 348, 242–245 (2015).

66. Foote, A. D. et al. Genome-culture coevolution promotes rapid divergence of killer whale ecotypes. Nat. Commun. 7, 11693 (2016).

67. Gonzalez, E. et al. The influence of CCL3L1 gene-containing segmental duplications on HIV-1/AIDS susceptibility. Science (80-.). 307, 1434–1440 (2005).

68. Johnson, M. E. et al. Positive selection of a gene family during the emergence of humans and African apes. Nature 413, 514–519 (2001).

69. Redon, R. et al. Global variation in copy number in the human genome. Nature 444, 444–454 (2006).

70. Paudel, Y. et al. Evolutionary dynamics of copy number variation in pig genomes in the context of adaptation and domestication. BMC Genomics 14, (2013).

71. Gao, Y. et al. CNV discovery for milk composition traits in dairy cattle using whole genome resequencing. BMC Genomics 18, (2017).

72. Zhang, R. Q., Wang, J. J., Zhang, T., Zhai, H. L. & Shen, W. Copy-number variation in goat genome sequence: A comparative analysis of the different litter size trait groups. Gene 696, 40–46 (2019).

73. Chen, C. et al. A comprehensive survey of copy number variation in 18 diverse pig populations and identification of candidate copy number variable genes associated with complex traits. BMC Genomics 13, (2012).

74. Amsterdam, A., Dantes, A. & Liscovitch, M. Role of phospholipase-D and phosphatidic acid in mediating gonadotropin-releasing hormone-induced inhibition of preantral granulosa cell differentiation. Endocrinology 135, 1205–1211 (1994).

75. Adhikari, D. et al. Tsc/mTORC1 signaling in oocytes governs the quiescence and activation of primordial follicles. Hum. Mol. Genet. 19, 397–410 (2009).

76. Tuppi, M. et al. Oocyte DNA damage quality control requires consecutive interplay of CHK2 and CK1 to activate p63. Nat. Struct. Mol. Biol. 25, 261–269 (2018).

77. Böing, M., Brand-Saberi, B. & Napirei, M. Murine transcription factor Math6 is a regulator of placenta development. Sci. Rep. 8, 14997 (2018).

78. Qiu, Y. et al. Carboxyl ester lipase is highly conserved in utilizing maternal supplied lipids during early development of zebrafish and human. Biochim. Biophys. Acta - Mol. Cell Biol. Lipids 1865, 158663 (2020).

79. Miller, R. & Lowe, M. E. Carboxyl ester lipase from either mother’s milk or the pancreas is required for efficient dietary triglyceride digestion in suckling mice. J. Nutr. 138, 927–930 (2008).

80. Wei, J. et al. An Integrative Genomic Analysis of the Superior Fecundity Phenotype in QSi5 Mice. Mol. Biotechnol. 53, 217–226 (2013).

81. Kosova, G., Scott, N. M., Niederberger, C., Prins, G. S. & Ober, C. Genome-wide association study identifies candidate genes for male fertility traits in humans. Am. J. Hum. Genet. 90, 950–961 (2012).

82. Coster, A. et al. The imprinted gene DIO3 is a candidate gene for litter size in pigs. PLoS One 7, e31825 (2012).

83. Magee, D. A. et al. Single nucleotide polymorphisms within the bovine DLK1-DIO3 imprinted domain are associated with economically important production traits in cattle. J. Hered. 102, 94–101 (2011).

84. Tao, L. et al. Combined approaches to reveal genes associated with litter size in Yunshang black goats. Anim. Genet. 51, 924–934 (2020).

85. Morgan, K., Harr, B., White, M. A., Payseur, B. A. & Turner, L. M. Disrupted gene networks in subfertile hybrid house mice. Mol. Biol. Evol. 37, 1547–1562 (2020).

86. Flegel, C. et al. Characterization of the Olfactory Receptors Expressed in Human Spermatozoa. Front. Mol. Biosci. 2, 73 (2016).

87. Daei-Farshbaf, N. et al. Expression pattern of olfactory receptor genes in human cumulus cells as an indicator for competent oocyte selection. Turkish J. Biol. 44, 371–380 (2020).

88. Langhammer, M. et al. High-fertility phenotypes: two outbred mouse models exhibit substantially different molecular and physiological strategies warranting improved fertility. Reproduction 147, 427–433 (2014).

89. Zeberg, H., Kelso, J. & Pääbo, S. The Neandertal Progesterone Receptor. Mol. Biol. Evol. 37, 2655–2660 (2020).

90. Lv, X. et al. Timely expression and activation of YAP1 in granulosa cells is essential for ovarian follicle development. FASEB J. 33, 10049–10064 (2019).

91. Anand-Ivell, R. & Ivell, R. Regulation of the reproductive cycle and early pregnancy by relaxin family peptides. Mol. Cell. Endocrinol. 382, 472–479 (2014).

92. Lei, W. et al. Progesterone and DNA damage encourage uterine cell proliferation and decidualization through up-regulating ribonucleotide reductase 2 expression during early pregnancy in mice. J. Biol. Chem. 287, 15174–15192 (2012).

93. Bünger, L., Renne, U., Dietl, G. & Kuhla, S. Long-term selection for protein amount over 70 generations in mice. Genet. Res. 72, 93–109 (1998).

94. Tsuneki, H., Wada, T. & Sasaoka, T. Role of orexin in the regulation of glucose homeostasis. Acta Physiol. 198, 335–348 (2010).

95. Zhang, Y. et al. QTL-based association analyses reveal novel genes influencing pleiotropy of metabolic syndrome (MetS). Obesity 21, 2099–2111 (2013).

96. Taussat, S. et al. Gene networks for three feed efficiency criteria reveal shared and specific biological processes. Genet. Sel. Evol. 52, 1–14 (2020).

97. Liu, B. & Mao, N. Smad5: Signaling roles in hematopoiesis and osteogenesis. Int. J. Biochem. Cell Biol. 36, 766–770 (2004).

98. Taye, M. et al. Deciphering signature of selection affecting beef quality traits in Angus cattle. Genes and Genomics 40, 63–75 (2018).

99. Jiao, S., Maltecca, C., Gray, K. A. & Cassady, J. P. Feed intake, average daily gain, feed efficiency, and real-time ultrasound traits in Duroc pigs: II. Genomewide association. J. Anim. Sci. 92, 2846–2860 (2014).

100. Xu, H. et al. A deletion downstream of the CHCHD7 gene is associated with growth traits in sheep. Animals 10, 1–10 (2020).

101. An, B. et al. Genome-wide association study reveals candidate genes associated with body measurement traits in Chinese Wagyu beef cattle. Anim. Genet. 50, 386–390 (2019).

102. Schrauwen, I. et al. FAM92A Underlies Nonsyndromic Postaxial Polydactyly in Humans and an Abnormal Limb and Digit Skeletal Phenotype in Mice. J. Bone Miner. Res. 34, 375–386 (2019).

103. Tsuchiya, M. et al. Cell surface flip-flop of phosphatidylserine is critical for PIEZO1-mediated myotube formation. Nat. Commun. 9, 1–15 (2018).

104. Rode, B. et al. Piezo1 channels sense whole body physical activity to reset cardiovascular homeostasis and enhance performance. Nat. Commun. 8, 1–11 (2017).

105. Göddeke, S. et al. CDH13 abundance interferes with adipocyte differentiation and is a novel biomarker for adipose tissue health. Int. J. Obes. 42, 1039–1050 (2018).

106. Teng, M. S., Wu, S., Hsu, L. A., Chou, H. H. & Ko, Y. L. Differential Associations between CDH13 Genotypes, Adiponectin Levels, and Circulating Levels of Cellular Adhesive Molecules. Mediators Inflamm. 2015, (2015).

107. Philippova, M. et al. A guide and guard: The many faces of T-cadherin. Cell. Signal. 21, 1035–1044 (2009).

108. Brenmoehl, J. et al. Browning of subcutaneous fat and higher surface temperature in response to phenotype selection for advanced endurance exercise performance in male DUhTP mice. J. Comp. Physiol. B Biochem. Syst. Environ. Physiol. 187, 361–373 (2017).

109. Lin, J. C., Chi, Y. L., Peng, H. Y. & Lu, Y. H. RBM4–Nova1–SRSF6 splicing cascade modulates the development of brown adipocytes. Biochim. Biophys. Acta - Gene Regul. Mech. 1859, 1368–1379 (2016).

110. Keller, M. A. et al. A gatekeeper helix determines the substrate specificity of Sjögren-Larsson Syndrome enzyme fatty aldehyde dehydrogenase. Nat. Commun. 5, 1–12 (2014).

111. Loro, E. et al. Effect of Interleukin-15 Receptor Alpha Ablation on the Metabolic Responses to Moderate Exercise Simulated by in vivo Isometric Muscle Contractions. Front. Physiol. 10, 1439 (2019).

112. Jiao, H. et al. Association analysis of positional obesity candidate genes based on integrated data from transcriptomics and linkage analysis. Int. J. Obes. 32, 816–825 (2008).

113. Duran, J. et al. Overexpression of ubiquitous 6-phosphofructo-2-kinase in the liver of transgenic mice results in weight gain. Biochem. Biophys. Res. Commun. 365, 291–297 (2008).

114. Sagara, S. et al. Overexpression of coupling factor 6 attenuates exercise-induced physiological cardiac hypertrophy by inhibiting PI3K/Akt signaling in mice. J. Hypertens. 30, 778–786 (2012).

