## supplementary information for "Genomic characterization of world’s longest selection experiment in mouse reveals the complexity of polygenic traits"

### Supplementary Material

#### 1. Establishment of the Dummerstorf mouse lines

During the years 1969 and 1970, four outbred strains (NMRI orig., Han:NMRI, CFW, CF1) and four inbred strains (CBA/Bln, AB/Bln, C57BL/Bln, XVII/Bln) were systematically crossed to establish the line FZTDU (*Forschungszentrum für Tierproduktion Dummerstorf*) [1,2]. Full-sib mating was avoided by random mating. This line has been kept unselected for almost 200 generations and originally maintained with 200 breeding pairs per generation until animals were relocated on generation ~160 and the number of breeding pairs dropped to 110. Thereafter, the number of breeding pairs has been kept at 125 (the current number of breeding pairs) avoiding consanguinity by random mating. All the Dummerstorf selection lines were derived from FZTDU starting at different time-points.

The process of selection to establish the fertility lines DUK and DUC began shortly after the establishment of FZTDU. In 1971, each fertility line was started with 60 breeding pairs. These animals were drawn from FZTDU by phenotypic maternal selection for number of offspring and litter weight at birth in the first litter (for more details see [3]). Selection for fertility continues to this day spanning more than 190 generations. The lowest number of breeding pairs was 19 (DUK) and 24 (DUC) at generation ~164 when animals were transferred to a facility by embryo transfer. Nowadays, the number of breeding pairs is 60 for both lines. Before relocation, the number of breeding pairs per generation and the selection intensity ranged between 60% and 100% and between 25% and 45%, respectively.

The body mass line DU6 was started in 1975 by phenotypic selection of FZTDU mice for high male body mass at 42 days of age. DU6 was founded by mating 80 pairs at around 9 weeks of age. The number of breeding pairs until generation ~154 was kept at 60-100 pairs. On generation ~154, animals were transferred to the new facility and the number of breeding pairs decreased to only 7 because of embryo transfer yield. Thereafter the number of breeding pairs was increased to 60. The selection strategy was later changed on generation 161 to selection based on estimated breeding values.

The breeding values for body mass at day 42 were calculated using mixed model with BLUP (Best Linear Unbiased Prediction) method [4]. As of generation 173 the proportion of female to male breeders has been kept at 2:1 (120 females and 60 males approx.) to mitigate the decreased pregnancy rate observed in DU6 females. This line continues to be selected (selection intensity of 45-90%). Also in 1975, the DU6P line was established by selection for high protein amount in male carcass at 42 days of age. The number of breeding pairs at the start of this line was 80 and was kept at 60-80 breeding pairs per generation. Then, on generation ~154, animals were relocated with 19 breeding pairs as founders, which were then increased to 60 pairs. Selection in DU6P stopped at generation 152 and the line is currently preserved without selection pressure by allowing random mating.

Finally, the high endurance line DUhLB was started in 1982 based on selection for high treadmill performance. It is thus the youngest of the Dummerstorf selection lines, as well as the shortest selected one (selection stopped at generation 141). Male running performance was evaluated based on distance (meters) covered on a treadmill before exhaustion (submaximal test).Trials were conducted after mating at 11 weeks of age. Subjects had no previous access to any kind of equipment that would influence their performance (untrained). Offspring of the highest scoring subjects were chosen for breeding. The line was founded with 100 breeding pairs and a selection intensity of 40% for the first 25 generations. Thereafter, the line was maintained with 60-80 breeding pairs at a selection intensity of 45-100%. On generation ~120, 44 breeding pairs were used as founders after transferring to the new facility. Together with DU6P, DUhLB is currently preserved without selection.

As indicated above, on generation 120-164 (year 2011 and 2012) a significant demographic event took place when all Dummerstorf mouse lines were moved from a conventional semi-barrier housing into a specific pathogen-free (SPF) environment. This transition was done via embryo transfer, which did not allow relocating entire populations. Therefore, lines were reestablished after a dramatic reduction in population size.

#### 2. Structural Variant Calling

Mapped and deduplicated short PE reads were used in detection of structural variants. As depth of coverage of reads mapped to the reference mouse genome sequence varied between 5 and 20x, we have split samples for each line into a high (10 samples) and low coverage (15 samples) set, and conducted structural variation analysis separately on these sets.

Three SV callers, Manta v.1.6.0[5], Whamg v.1.7.0[6] and Lumpy v.0.2.13[7] were selected based on their sensitivity and precision[8]. Manta integrates paired-read (PR), split-read (SR) evidence and SV breakend assembly (AS) during SV discovery. Whamg implements PR and SR support while Lumpy relies on a probabilistic copy number variation discovery combing PR, SR and read-depth (RD). Intersecting results of multiple SV callers applying different SV detection approaches has previously been shown to improve accuracy of variant call sets[8,9].

Manta SV calls in genomic regions with depth greater than 3x the median chromosome depth near one or both SV breakends mainly caused by reads mapping in low complexity regions, as well as reads with MAPQ<30 were filtered out. Furthermore, variant calls for which samples did not pass Manta caller quality filters, and have genotype quality below 20 (GQ<20) have been filtered out. Calls with paired-read (PR) and split-read (SR) support of PR>=3 and SR>=3 were retained.

Whamg SV calls of size <50bp and >2Mb were filtered out, along with calls with fewer than 5 supporting reads and GQ<20. Calls associated with poorly mapped regions and BND-type calls with high cross-chromosomal mapping scores (CW>0.2) were removed, as Whamg does not specifically call translocations. Lumpy calls for which evidence supporting the variant were below 5 (SU<5) and calls with GQ<20 were filtered out. Both Whamg and Lumpy SV call sets were genotyped with Svtyper v0.7.1[10].

Unlocalized and unplaced scaffolds have been removed from all SV sets and only scaffolds assigned to chromosomes have been included in further analysis. Survivor v.1.0.7[11] was used to merge SV call sets within and among samples. For each mice line sample, we first merged SV events of the same type, called by at least two SV callers, with start/ end positions detected within +/-1000 bp, identified in at least 60% of samples in a low coverage set (10 samples out of 15) and 100% of samples in a high coverage set (all 10 samples) for each mice line.

The union of SVs detected in two separate sample sets for each line were further used. We then intersected SV calls among all mice lines to obtain SVs private for each mice line (line-specific) and shared among lines (Supplementary Figure 8). To further reduce the FDR, SV calls overlapping gaps and high coverage regions (> 80x) in the reference genome assembly were filtered out. High coverage regions were determined for each mouse line based on the intersection among samples of 1-kb windows containing reads mapping with depth of coverage > 80x. Variants specified as “BND” (translocations) were removed and deletions, and duplications and inversions were further investigated (Supplementary Figure 9-12).

We annotated the final SV set with the Ensembl VEP v. 101.0[12] focusing on variants overlapping protein-coding genes (maximum SV size = 200 Mb). Functional classification of genes was based on literature and database search (OrthoDB v10[13]; Uniprot[14]; NCBI Entrez gene[15]), and Gene Ontology enrichment analysis (Shiny GO, FDR < 0.05[16]).

### Supplementary figures

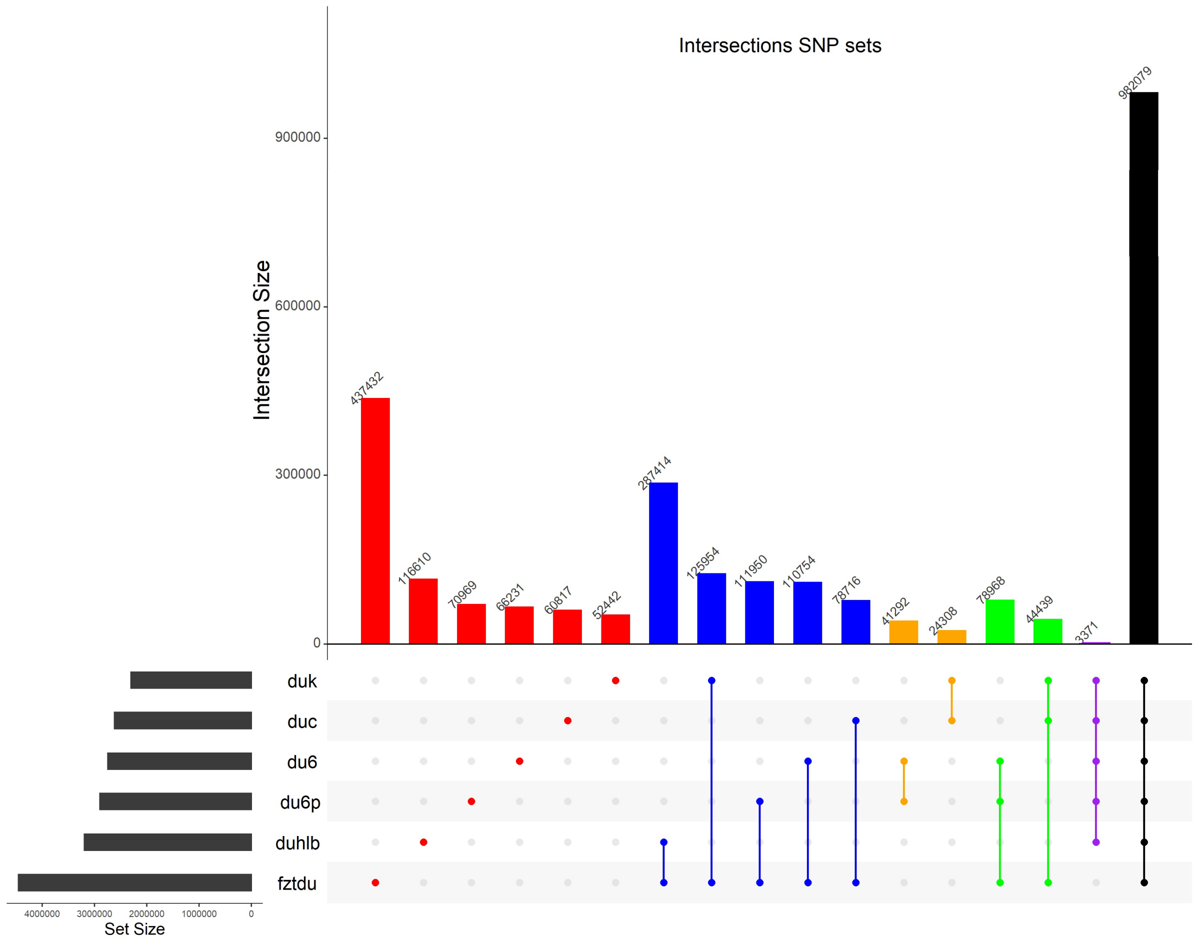


**Supplementary Figure 1. Number of private and shared SNPs among lines.** Each bar corresponds to the number of SNPs that are private for each line or shared by two or more lines. For practical purposes, the intersections shown correspond to a subset of all possible intersection among the 6 lines.

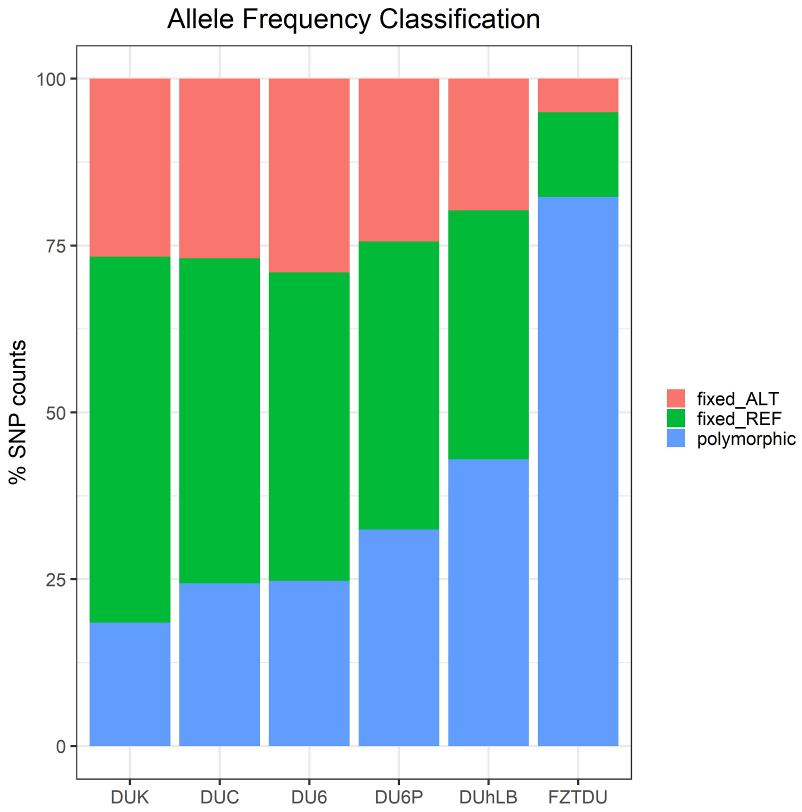


**Supplementary Figure 2. SNP allele frequency state classification**. Sites in which SNPs were observed across the whole set of samples are classified for each mouse line as fixed-alternative (alternative allele homozygous in all subjects, fixed_ALT), fixed-reference (reference allele homozygous in all subjects, fixed_REF) and polymorphic (not fixed).

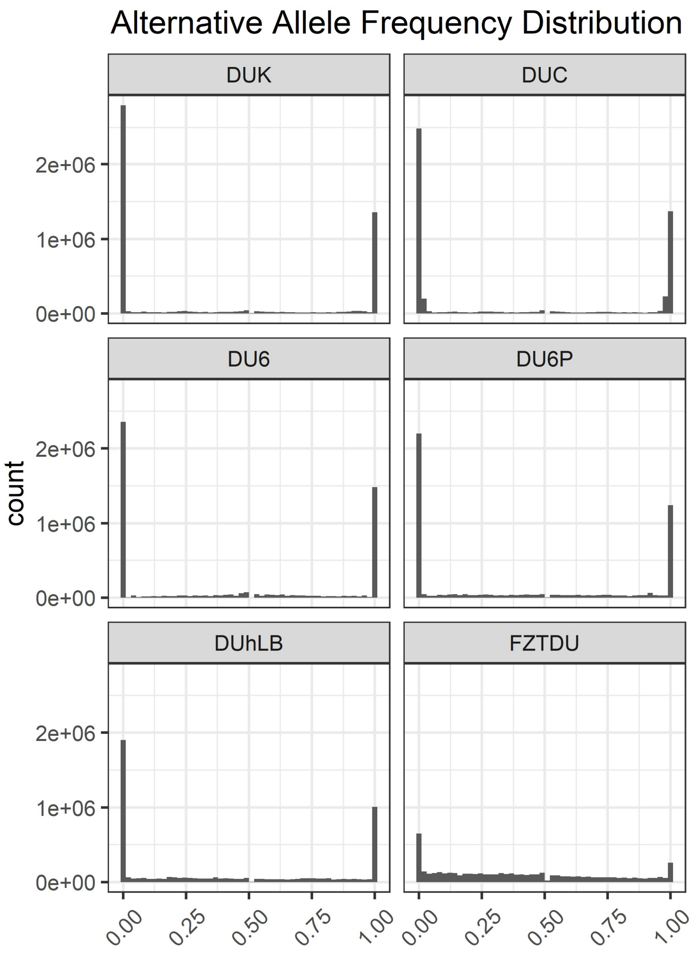


**Supplementary Figure 3. Alternative allele frequency distribution.** Counts of SNPs along the allele frequency spectrum for each mouse line.

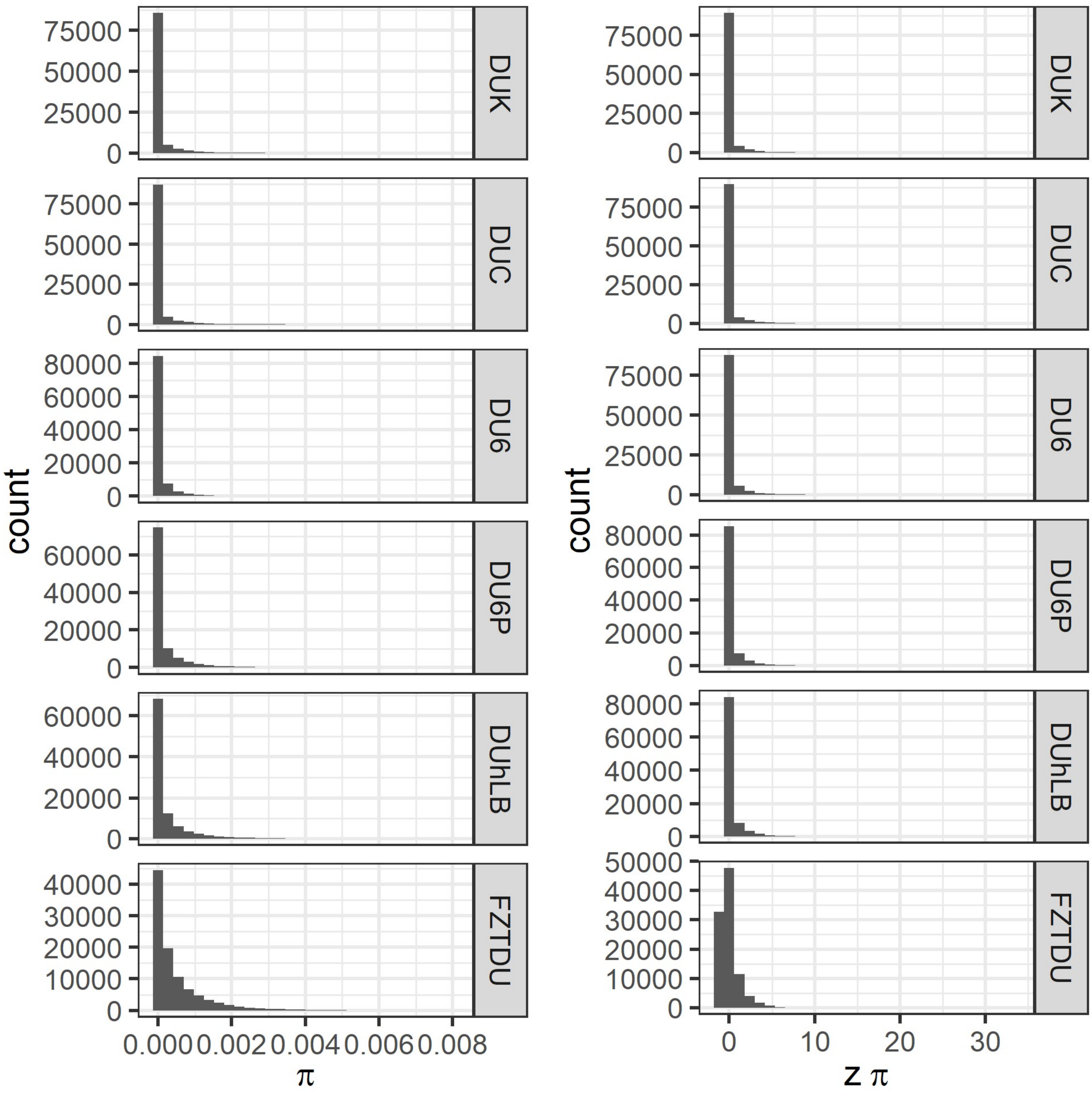


**Supplementary Figure 4. Nucleotide diversity (π) distribution in the Dummerstorf mouse lines.** Nucleotide diversity (π) was calculated in sliding window mode (size=50Kb, step=25Kb, ≥10 SNPs). Scores were transformed to z-scores in order to represent the data in terms of standard deviations from the genomic mean. The distributions illustrate the low levels of genetic diversity within lines, with the most scores accumulate at the lower end of the distribution (left), thus highly diverse regions are rare events.

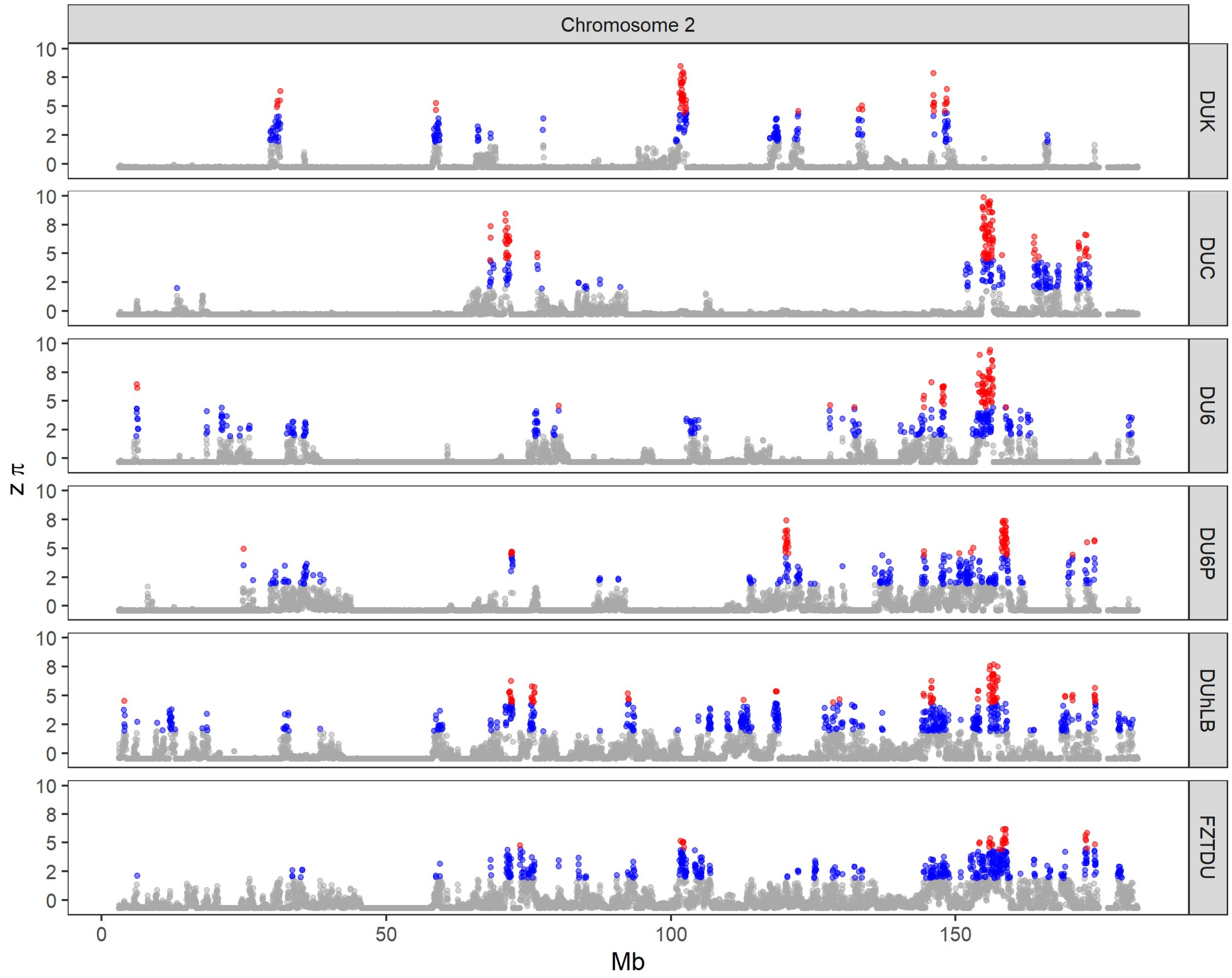


**Supplementary Figure 5. Example of one chromosome representative of the level of genetic diversity observed in the Dummerstorf mouse lines.** The stretches of low diversity are longer and more abundant than in FZTDU. Regions of extreme genetic diversity are shown in blue (top 5% most diverse windows) and red (top 1% most diverse windows).

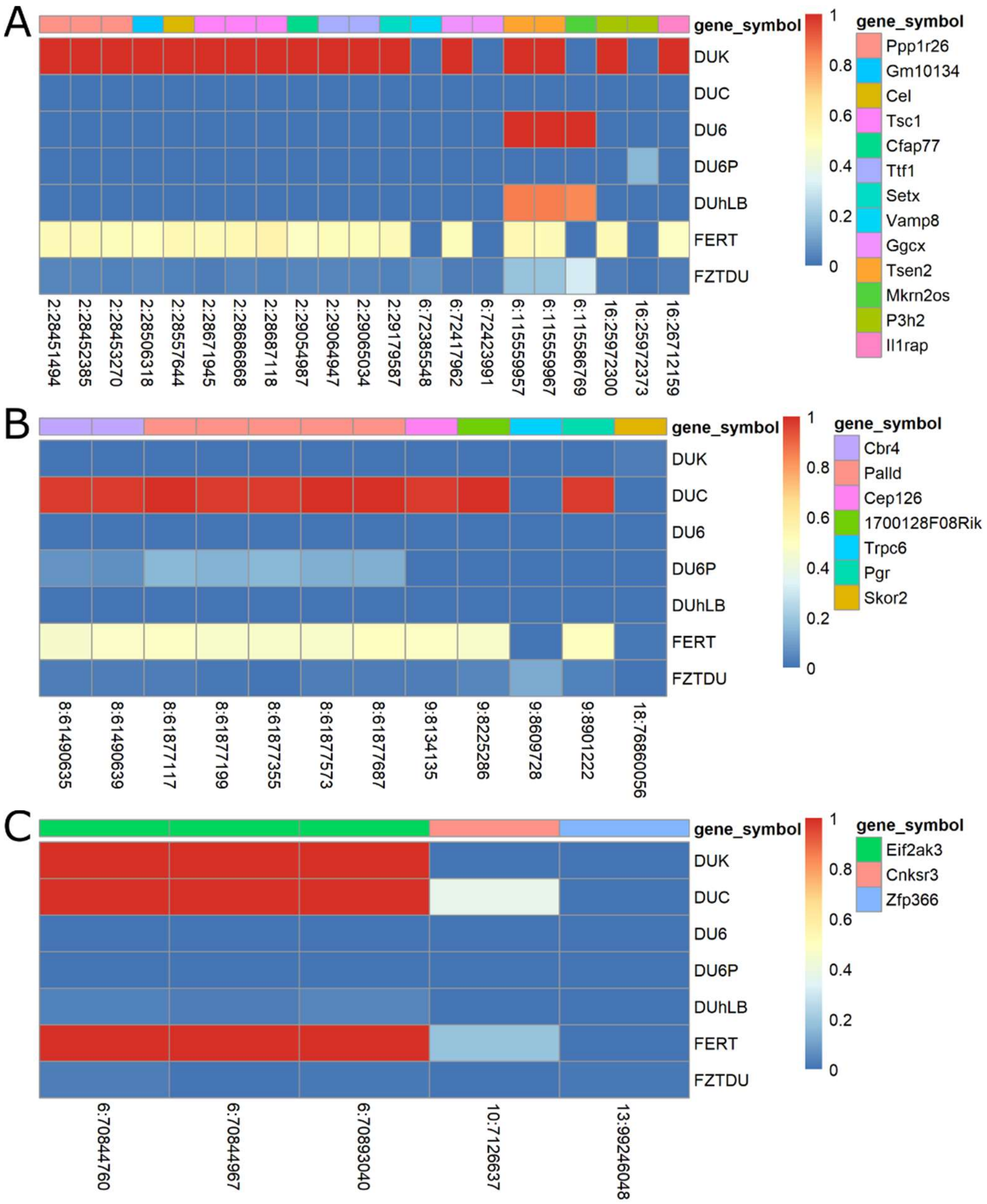


**Supplementary Figure 6. Allele frequency heatmap of non-synonymous mutations in RDD genes.**

Allele frequencies of non-synonymous SNPs in genes overlapping regions of distinct genetic differentiation for DUK (**A**), DUC (**B**) and the joint fertility population FERT (**C**). The gradient scale represents the allele frequency from low (blue) to high (red).

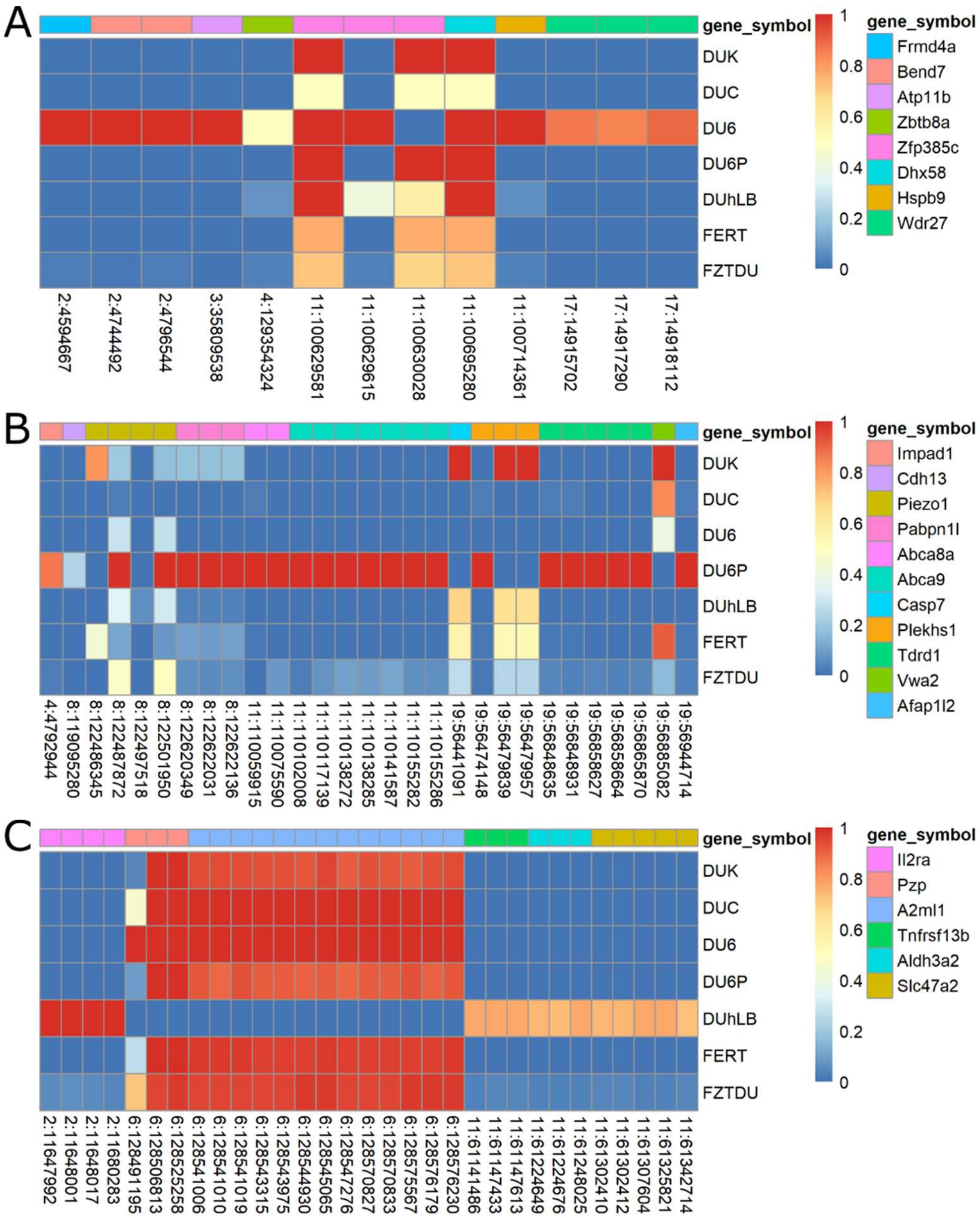


**Supplementary Figure 7. Allele frequency heatmap of non-synonymous mutations in RDD genes.** Allele frequencies of non-synonymous SNPs in genes overlapping regions of distinct genetic differentiation for DU6 (**A**), DU6P (**B**) and DUhLB (**C**). The gradient scale represents the allele frequency from low (blue) to high (red).

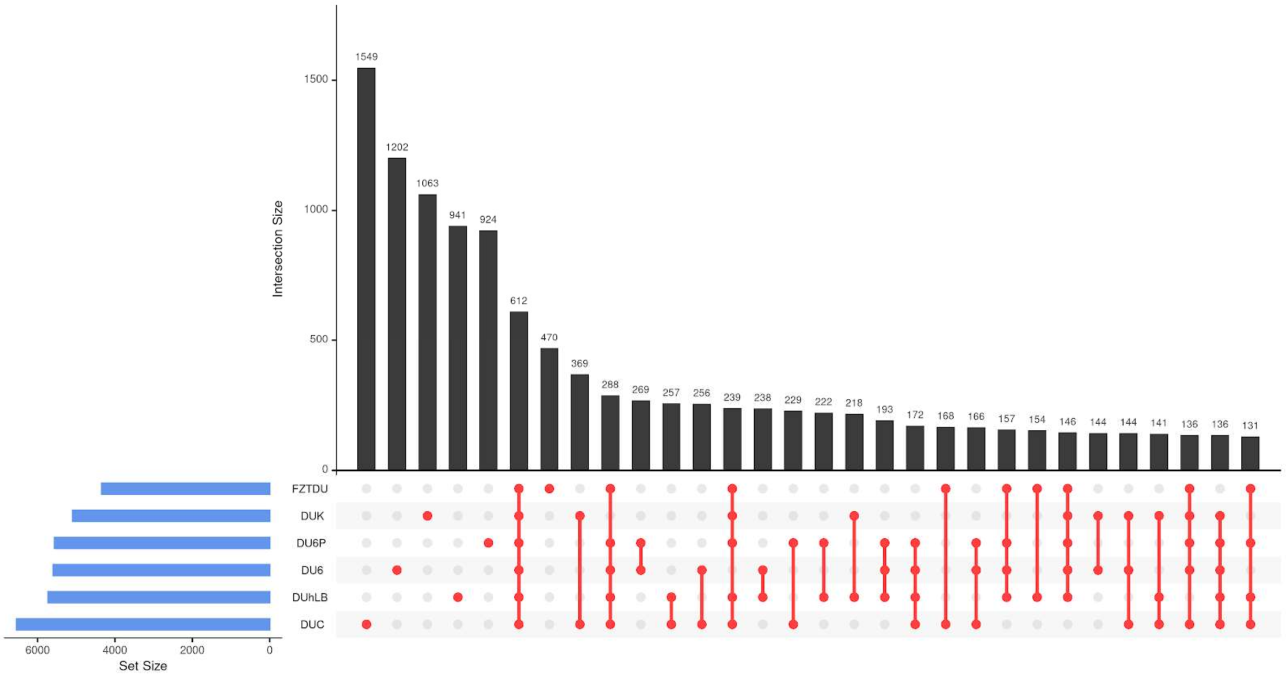


**Supplementary Figure 8. Shared and line-specific structural variants.** SVs detected in union of high and low coverage sample sets for each mice line.

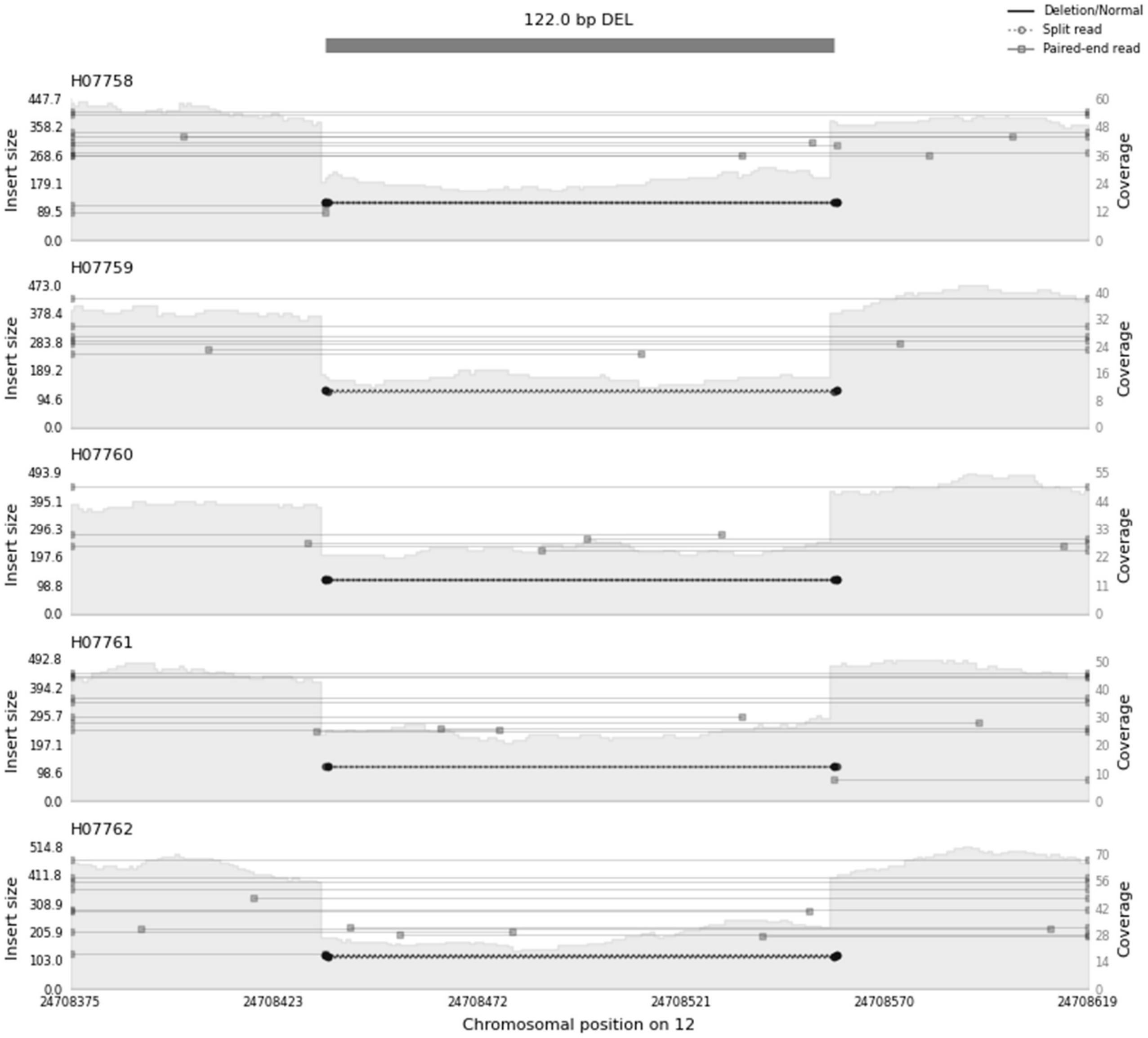


**Supplementary Figure 9. Example of a polymorphic deletion.** SV is shown in 5 out of 10 samples of the high coverage set, indicating the genomic location (x-axis), the insert size (y-axis, left) and coverage (y-axis-right).

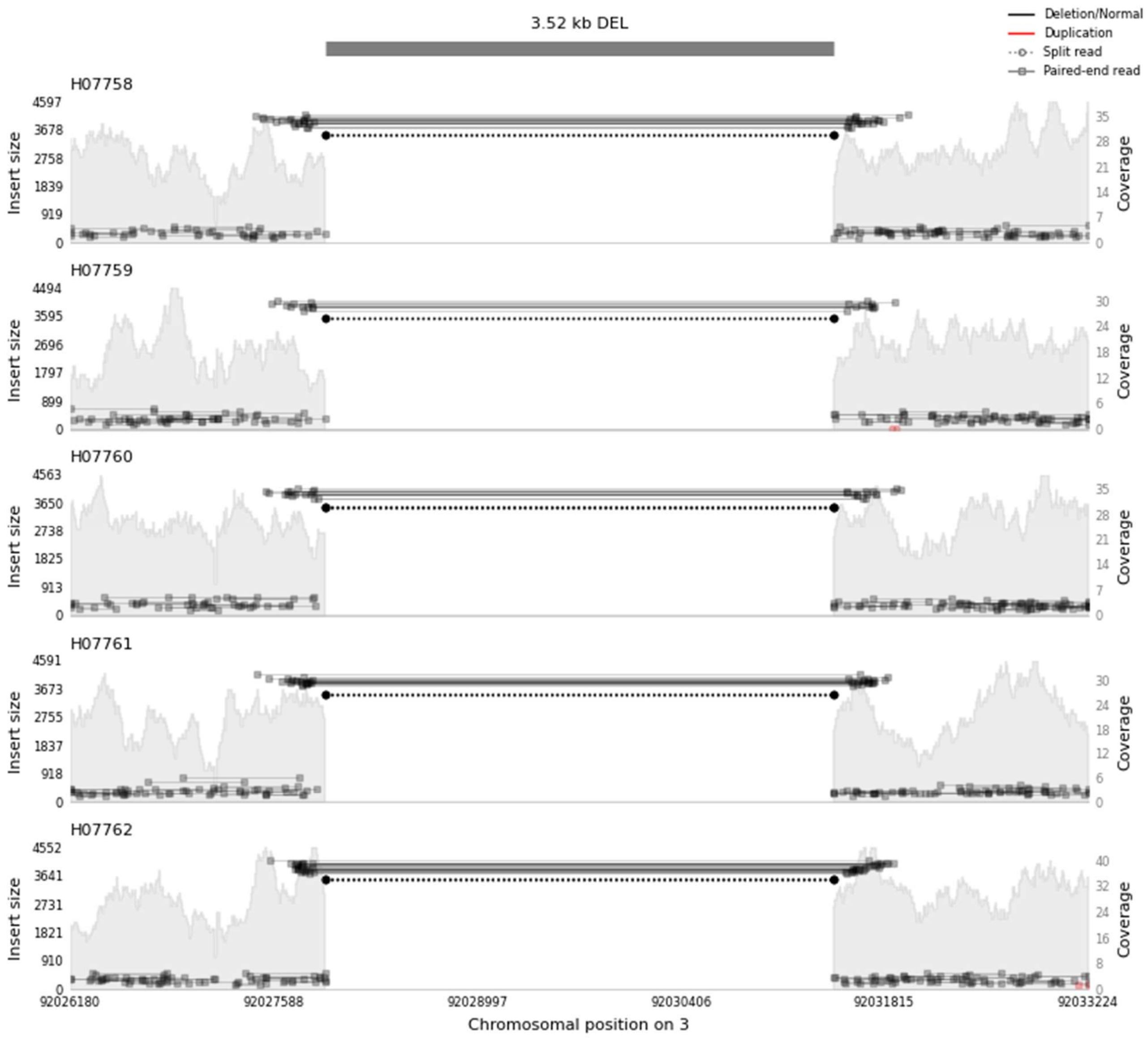


**Supplementary Figure 10. Example of a fixed deletion.** SV is shown in 5 out of 10 samples of the high coverage set, indicating the genomic location (x-axis), the insert size (y-axis, left) and coverage (y-axis-right).

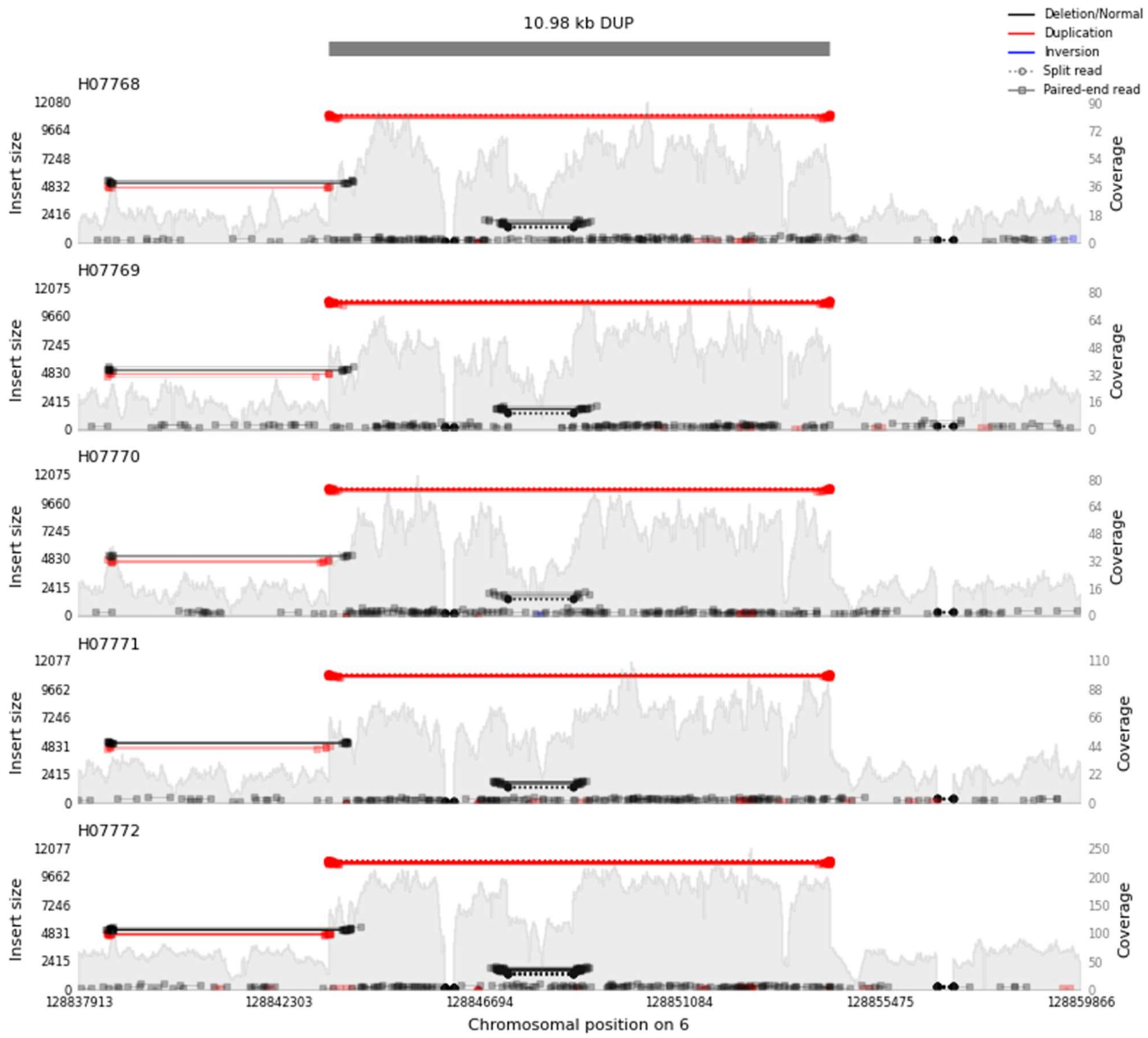


**Supplementary Figure 11. Example of a fixed duplication.** SV is shown in 5 out of 10 samples of the high coverage set, indicating the genomic location (x-axis), the insert size (y-axis, left) and coverage (y-axis-right).

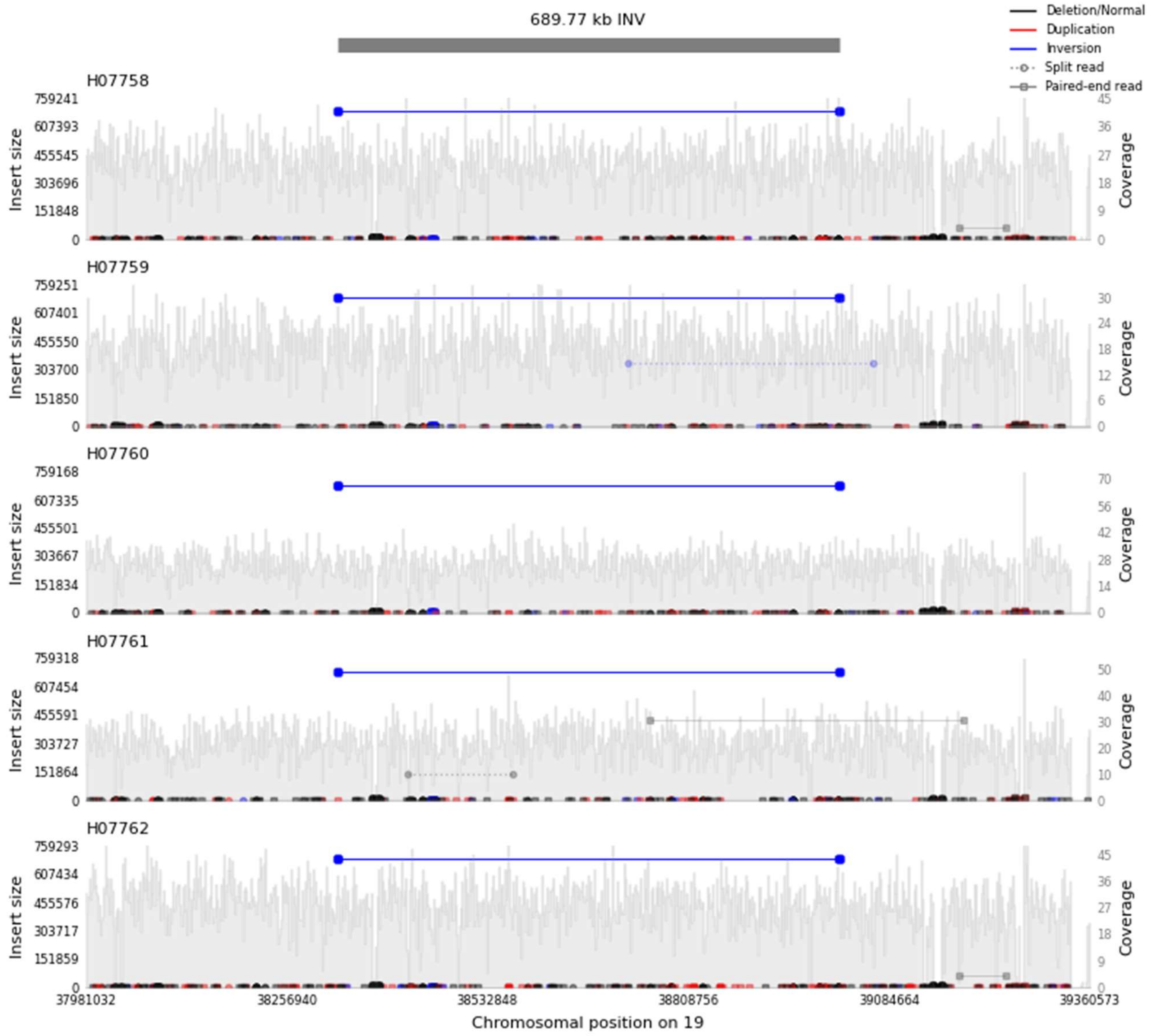


**Supplementary Figure 12. Example of a fixed inversion.** SV is shown in 5 out of 10 samples of the high coverage set, indicating the genomic location (x-axis), the insert size (y-axis, left) and coverage (y-axis-right).

### Supplementary tables

**Supplementary Table 1. Number of private variants with predicted high/moderate effects according to SnpEff.**

| | **SNPs** | **INDELs** | **Genes** |
|---|---|---|---|
| DUK | 996 | 142 | 557 |
| DUC | 640 | 147 | 509 |
| DU6 | 752 | 167 | 582 |
| DU6P | 783 | 153 | 576 |
| DUhLB | 1970 | 231 | 1072 |

Private variants for FZTDU not included as this line was unselected.

**Supplementary Table 2. Significantly enriched terms based on RDD gene lists**

| Line | **Term/Pathway** | **Genes** | **FDR** |
|---|---|---|---|
| DUK | mmu04072: Phospholipase D signaling pathway | Raf1 | 0.0082875 |
| | | Adcy6 | |
| | | Grm8 | |
| | | Tsc1 | |
| | | Ralgds | |
| DUC | GO:0030518: intracellular steroid hormone receptor signaling pathway | Pias2 | 0.030545 |
| | | Pgr | |
| | | Rxfp1 | |
| | | Yap1 | |
| DUC | GO:0048814: regulation of dendrite morphogenesis | Pias2 | 0.074441 |
| | | Trpc6 | |
| | | Skor2 | |
| DUC | GO:0030518: intracellular steroid hormone receptor signaling pathway | Pias2 | 0.074441 |
| | | Pgr | |
| | | Yap1 | |
| DUhLB | mmu00340: Histidine metabolism | Aldh3a1 | 0.099122 |
| | | Aldh3a2 | |
| DUhLB | mmu00410: beta-Alanine metabolism | Aldh3a1 | 0.099122 |
| | | Aldh3a2 | |

Significantly enriched GO terms and pathways at FDR < 0.1

**Supplementary Table 3. Proportion of line-specific fixed and polymorphic structural variants in genic regions**

| | Fixed | | Polymorphic | |
|---|---|---|---|---|
| | Number | Length (mean) | Number | Length (mean) |
| DUK | 5 | 13.58 (2.7) kbp | 15 | 204.62 (13.64) Mb |
| DUC | 2 | 2.19 kbp (1.1 kbp) | 34 | 186.92 (5.5) Mb |
| DU6 | 4 | 7.75 (1.94) kbp | 14 | 302 (21.57) kb |
| DU6P | 7 | 29.11 (4.65) kbp | 17 | 129.96 (7.6) Mb |
| DUhLB | 6 | 11.16 (1.9) kbp | 14 | 3.72 (0.265) Mb |
| FZTDU | 1 | 1.14 kbp | 8 | 14.09 (1.8) Mb |

**Supplementary Table 4. Types and lengths of line-specific fixed structural variants in genic regions**

| | DEL | DEL length | DUP | DUP length | INV | INV length |
|---|---|---|---|---|---|---|
| DUK | 5 | 13.6 Kb | -- | -- | -- | -- |
| DUC | 1 | 1.3 Kb | -- | -- | 1 | 0.923 Kb |
| DU6 | 4 | 7.7 Kb | -- | -- | -- | -- |
| DU6P | 3 | 8.5 Kb | 1 | 11 Kb | 3 | 9.6 Kb |
| DUhLB | 4 | 3.3 Kb | -- | -- | 2 | 7.9 Kb |
| FZTDU | 1 | 1.1 Kb | -- | -- | -- | -- |

**Supplementary Table 5. Number of genes affected by line-specific fixed and polymorphic structural variants**

| | Fixed | | | Polymorphic | | |
|---|---|---|---|---|---|---|
| | DEL | DUP | INV | DEL | DUP | INV |
| DUK | 5 | -- | -- | 4 | 28 | 1694 |
| DUC | 1 | -- | 1 | 9 | 38 | 1363 |
| DU6 | 4 | -- | -- | 6 | -- | 7 |
| DU6P | 3 | 1 | 3 | 11 | -- | 1130 |
| DUhLB | 4 | -- | 2 | 11 | 3 | 7 |
| FZTDU | 1 | -- | -- | 3 | -- | 266 |

**Supplementary Table 6. Number of genes in functional groups affected by line-specific structural variants**

| | DUK | DUC | DU6 | DU6P | DUhLB | FZTDU |
|---|---|---|---|---|---|---|
| **Reproduction** | 8 | 13 | 1 | 1 | 3 | 1 |
| **Metabolism/Energy conversion** | -- | 27 | 3 | 16 | 8 | -- |
| **Immune system** | -- | 3 | 1 | 4 | 1 | 13 |
| **Nervous system** | 5 | 13 | 1 | 3 | 4 | 1 |
| **Cardiovascular system** | 2 | 1 | 1 | 1 | 2 | 15 |
| **Endocrine system** | 2 | 2 | 2 | 1 | 2 | -- |
| **Sensory perception** | 297 | 36 | 2 | -- | 3 | -- |
| **Other (cell cycle, transcription)** | 2 | 88 | 3 | 15 | 1 | 2 |

#### Supplementary references


1. Dietl, G., Langhammer, M. & Renne, U. Model simulations for genetic random drift in the outbred strain Fzt: DU. *Arch. FUR TIERZUCHT* **47**, 595–604 (2004).
2. Schueler, L. Mouse strain Fzt:DU and its use as model in animal breeding research. *Arch. für Tierzucht (Archives Anim. Breeding)* **28**, 357–363 (1985).
3. Langhammer, M. *et al.* High-fertility phenotypes: two outbred mouse models exhibit substantially different molecular and physiological strategies warranting improved fertility. *Reproduction* **147**, 427–433 (2014).
4. Henderson, C. R. Best Linear Unbiased Estimation and Prediction under a Selection Model. *Biometrics* **31**, 423–447 (1975).
5. Chen, X. *et al.* Manta: Rapid detection of structural variants and indels for germline and cancer sequencing applications. *Bioinformatics* **32**, 1220–1222 (2016).
6. Kronenberg, Z. N. *et al.* Wham: Identifying Structural Variants of Biological Consequence. *PLOS Comput. Biol.* **11**, e1004572 (2015).
7. Layer, R. M., Chiang, C., Quinlan, A. R. & Hall, I. M. LUMPY: A probabilistic framework for structural variant discovery. *Genome Biol.* **15**, R84 (2014).
8. Kosugi, S. *et al.* Comprehensive evaluation of structural variation detection algorithms for whole genome sequencing. *Genome Biol.* **20**, (2019).
9. Pirooznia, M., Goes, F. & Zandi, P. P. Whole-genome CNV analysis: Advances in computational approaches. *Front. Genet.* **6**, (2015).
10. Chiang, C. *et al.* SpeedSeq: Ultra-fast personal genome analysis and interpretation. *Nat. Methods* **12**, 966–968 (2015).
11. Jeffares, D. C. *et al.* Transient structural variations have strong effects on quantitative traits and reproductive isolation in fission yeast. *Nat. Commun.* **8**, 1–11 (2017).
12. McLaren, W. *et al.* The Ensembl Variant Effect Predictor. *Genome Biol.* **17**, 122 (2016).
13. Kriventseva, E. V. *et al.* OrthoDB v10: Sampling the diversity of animal, plant, fungal, protist, bacterial and viral genomes for evolutionary and functional annotations of orthologs. *Nucleic Acids Res.* **47**, D807–D811 (2019).
14. Apweiler, R. *et al.* UniProt: The universal protein knowledgebase. *Nucleic Acids Res.* **32**, (2004).
15. Maglott, D., Ostell, J., Pruitt, K. D. & Tatusova, T. Entrez gene: Gene-centered information at NCBI. *Nucleic Acids Res.* **39**, D52–D57 (2011).
16. Ge, S. X., Jung, D., Jung, D. & Yao, R. ShinyGO: A graphical gene-set enrichment tool for animals and plants. *Bioinformatics* **36**, 2628–2629 (2020).
